# Chronic activation of the aryl hydrocarbon receptor in muscle exacerbates ischemic pathology in chronic kidney disease

**DOI:** 10.1101/2023.05.16.541060

**Authors:** Nicholas Balestrieri, Victoria Palzkill, Caroline Pass, Jianna Tan, Zachary R. Salyers, Chatick Moparthy, Ania Murillo, Kyoungrae Kim, Trace Thome, Qingping Yang, Kerri A. O’Malley, Scott A. Berceli, Feng Yue, Salvatore T. Scali, Leonardo F. Ferreira, Terence E. Ryan

## Abstract

Chronic kidney disease (CKD) accelerates the development of atherosclerosis, decreases muscle function, and increases the risk of amputation or death in patients with peripheral artery disease (PAD). However, the cellular and physiological mechanisms underlying this pathobiology are ill-defined. Recent work has indicated that tryptophan-derived uremic toxins, many of which are ligands for the aryl hydrocarbon receptor (AHR), are associated with adverse limb outcomes in PAD. We hypothesized that chronic AHR activation, driven by the accumulation of tryptophan-derived uremic metabolites, may mediate the myopathic condition in the presence of CKD and PAD. Both PAD patients with CKD and mice with CKD subjected to femoral artery ligation (FAL) displayed significantly higher mRNA expression of classical AHR-dependent genes (*Cyp1a1*, *Cyp1b1*, and *Aldh3a1*) when compared to either muscle from the PAD condition with normal renal function (*P*<0.05 for all three genes) or non-ischemic controls. Skeletal-muscle-specific AHR deletion in mice (AHR^mKO^) significantly improved limb muscle perfusion recovery and arteriogenesis, preserved vasculogenic paracrine signaling from myofibers, increased muscle mass and contractile function, as well as enhanced mitochondrial oxidative phosphorylation and respiratory capacity in an experimental model of PAD/CKD. Moreover, viral-mediated skeletal muscle-specific expression of a constitutively active AHR in mice with normal kidney function exacerbated the ischemic myopathy evidenced by smaller muscle masses, reduced contractile function, histopathology, altered vasculogenic signaling, and lower mitochondrial respiratory function. These findings establish chronic AHR activation in muscle as a pivotal regulator of the ischemic limb pathology in PAD. Further, the totality of the results provide support for testing of clinical interventions that diminish AHR signaling in these conditions.

## INTRODUCTION

Peripheral arterial disease (PAD) affects 8-12 million Americans ^1^ and is the third highest leading cause of cardiovascular mortality ^2^. PAD is caused by atherosclerotic narrowing or occlusion in the lower extremities which leads to a spectrum of life-altering symptomatology including claudication, ischemic rest pain, as well as gangrene and/or non-healing ulcers that commonly result in major limb amputation especially when revascularization attempts fail (or are not possible). Complicating the etiology of PAD, patients typically present with one or more comorbid conditions or risk factors that accelerate disease evolution and are associated with poorer health outcomes. Among these, chronic kidney disease (CKD) is strongly associated with the development of atherosclerosis, decreased muscle function, and increased risk of amputation or death in PAD patients ^3–6^. For example, mortality risk for PAD patients with CKD is ∼2-to-4 times higher than PAD patients without CKD ^6–9^. CKD has also been shown to be associated with increased failure rates of both endovascular and open surgical revascularization procedures in PAD management ^3,10^. While the clinical evidence demonstrating that CKD exacerbates PAD pathobiology is undeniable, the cellular and physiological mechanisms underlying this pathology is ill-defined.

One function of the kidneys is to rid the body of waste materials that are either ingested or produced endogenously by normal metabolism. CKD leads to retention of waste products that can be harmful to other tissues/organs, including skeletal muscle ^11–15^ and blood vessels ^16–18^ where the combined effects presumably contribute to worsened PAD pathology. Unfortunately, many uremic metabolites are protein-bound and not adequately filtered by dialysis membranes and remain elevated despite treatment with renal replacement therapies ^19,20^. A recent report demonstrated a strong correlation between tryptophan-derived uremic metabolites and adverse limb events in PAD patients ^21^, highlighting the potential that specific biological pathways may contribute to the pathobiology of PAD in patients with CKD. Interestingly, many of these tryptophan-derived uremic metabolites have been shown to impair skeletal muscle mitochondrial function and drive atrophy ^22–25^, potentially implicating skeletal muscle function and mitochondrial health as a site of coalescence between pathobiology of CKD and PAD. Importantly, emerging evidence in PAD patients has demonstrated that muscle function/exercise capacity is a strong predictor of morbidity/mortality ^26–30^.

Interestingly, several established tryptophan-derived uremic metabolites (kynurenines and indoles) are ligands of the aryl hydrocarbon receptor (AHR), a ligand-activated transcription factor belonging to the basic helix-loop-helix/Per-ARNT-Sim (bHLH/PAS) family. Both mice and human patients with CKD display elevated expression of the *Ahr* ^31^. The most widely studied of the uremic metabolites, indoxyl sulfate, causes muscle atrophy and mitochondrial dysfunction in mice with normal kidney function ^22,23,32^, although the underlying mechanisms are unknown. Chronic activation of the AHR (or its downstream effectors *Cyp1a1* and *Cyp1b1*) has been shown to reduce angiogenesis ^33,34^ and increase atherosclerosis ^35–37^. A recent report demonstrated that chemical inhibition of the AHR could improve perfusion recovery in CKD mice following femoral artery ligation (FAL) and normalize post-ischemic angiogenesis ^21^. Comparatively, little is known about the role of the AHR in mediating the toxic effects of uremic metabolite accumulation in skeletal muscle exposed to the CKD condition. However, a recent study reported that AHR activation in skeletal muscle could phenocopy the effects of tobacco smoking, a major risk factor for PAD, on skeletal muscle pathology including atrophy and mitochondrial respiratory dysfunction ^38^. Further, strong associations between the levels of CKD-associated AHR-ligands and muscle mitochondrial energetics were recently reported in rodents with CKD ^25^. Based on the accumulation of evidence described above, we hypothesized that chronic AHR activation, driven by the accumulation of tryptophan-derived uremic metabolites, may mediate the myopathic condition in the presence of CKD and PAD.

## MATERIALS AND METHODS

### Human Subjects

Gastrocnemius muscle specimens were collected from non-PAD adults, PAD patients with normal renal function, and PAD patients with CKD via percutaneous muscle biopsy using sterile procedures as previously described^39,40^. This study was approved by the institutional review boards at the University of Florida and the Malcom Randall VA Medical Center (Protocol IRB201801553). All study procedures were carried out according to the Declaration of Helsinki and participants were fully informed about the research and informed consent was obtained.

### Animals

A skeletal muscle-specific AHR knockout mouse was generated by breeding floxed AHR mice (AHR^tm3.1Bra^/J, Jackson Laboratories, Stock No. 006203) with HSA-MCM mice that express MerCreMer double fusion protein under the control of the human *Acta1* (actin, alpha 1, skeletal muscle) promoter (Tg(ACTA1-cre/Esr1*)2Kesr/J, Jackson Laboratories, Stock No. 025750). All experiments involved male and female mice that were randomized to experimental groups such that both the surgeon and experimenters were blinded to the treatment groups and genotypes. Female mice underwent an ovariectomy one week prior to enrollment. For AAV experiments four-month-old male and ovariectomized female C57BL6J mice were purchased from Jackson Labs (Stock No. 000664). All animal experiments adhered to the *Guide for the Care and Use of Laboratory Animals* from the Institute for Laboratory Animal Research, National Research Council, Washington, D.C., National Academy Press. All procedures were approved by the Institutional Animal Care and Use Committee of the University of Florida (Protocol 202110484).

### AAV construction and delivery

To accomplish muscle cell-specific expression of transgenes, the human skeletal actin (*Acta1*; termed HSA herein) promoter (1541 bp proximal to the transcription start site) was PCR amplified from human genomic DNA isolated from a donor muscle biopsy. The AAV-HSA-GFP plasmid was developed by inserting a human HSA promoter and GFP (ZsGreen1) into the promoterless AAV vector (Cell BioLabs, Cat. No. VPK-411-DJ) using In-Fusion Cloning (Takara Bio, Cat. No. 638911). Similarly, the mouse *Ahr*, including the ligand binding domain, was PCR amplified from cDNA obtained from a C57BL6J mouse and inserted downstream of the HSA promoter. To generate a constitutively active *Ahr* (CAAHR) vector, the mouse AHR coding sequence was PCR amplified from cDNA obtained from a C57BL6J mouse such that the ligand binding domain (amino acids 277–418) was deleted, and subsequently inserted downstream of the HSA promoter. AAV9 was delivered via intramuscular injections to the surgical hind limb (gastrocnemius, tibialis anterior (TA), extensor digitorum longus (EDL)) muscles at a dosage of 5E+11 vg/limb.

### Induction of chronic kidney disease

All mice were randomized to either a casein control diet for 7 days followed by either a diet enriched with 0.2% adenine^41,42^ or maintained on the casein diet for three weeks until undergoing femoral artery ligation (FAL) surgery to induce limb ischemia. Following FAL surgery, the mice maintained their respective diet cohorts. Analysis of serum uremic toxin levels was carried out at the Southeast Center for Integrated Metabolomics at the University of Florida as previously described ^25,42^.

### Assessment of kidney function

Kidney function was evaluated via previously described glomerular filtration rate (GFR) and blood urea nitrogen (BUN) levels ^25,41–45^. Detailed methodology for these procedures can be found in the Supplemental Materials and Data.

### Animal model of peripheral artery disease

Femoral artery ligation (FAL) ^41^ was performed by anesthetizing mice with an intraperitoneal injection of ketamine (90 mg/kg) and xylazine (10 mg/kg) and surgically inducing unilateral hindlimb ischemia by placing silk ligatures on the femoral artery just distal the inguinal ligament and immediately proximal to the saphenous and popliteal branches. Sham surgeries involved the complete dissection and separation of the neurovascular bundle, but no ligation was performed. In all procedures, buprenorphine (0.05 mg/kg) was given post-operatively for analgesia.

### Limb perfusion assessment

Limb perfusion was assessed by laser Doppler flowmeter (moorVMS-LDF, Moor Instruments) prior to surgery, post-surgery day 3, 7, 10 post-surgery and just prior to sacrifice, as described previously ^41,46,47^. Perfusion recovery was calculated as a percentage of the non-ischemic limb.

### *Ex-vivo* muscle contractile function

As previously described, ^46,47^ the extensor digitorum longus (EDL) muscle was surgically removed and affixed to a dual-mode muscle lever system (Aurora Scientific; 300 C-LR) using silk sutures tightly tied to the proximal and distal tendons.

### Isolation of skeletal muscle mitochondria

Skeletal muscle mitochondria were isolated as previously described ^25,45^ from the gastrocnemius muscle. A detailed description of this methodology is provided in the Supplemental Materials and Data.

### Assessment of mitochondrial oxygen consumption

High-resolution respirometry was measured using Oroboros Oxygraph-2k (O2K) measuring oxygen flux (*J*O2) at 37°C in mitochondrial assay buffer (105 mM K-MES, 30 mM KCl, 1 mM EGTA, 10 mM K_2_HPO_4_, 5 mM MgCl_2_-6H_2_O, 2.5 mg/mL BSA, pH 7.2), supplemented with either 5 mM creatine (conductance assay) or 20 mM creatine (respiratory capacity assay). Detailed descriptions of the respirometery protocols can be found in the Supplemental Materials and Data.

### Skeletal muscle histology and immunofluorescence

Skeletal muscle histopathology was assessed in transverse sections of the tibialis anterior muscle using several approaches. Detailed methodology for each approach is provided in the Supplemental Materials and Data. Images displayed in figure panels were chosen to represent the average values of quantified outcome variables for each group. Validation of all immunofluorescence antibodies was performed using no primary control staining on parallel tissue sections.

### Immunoblotting

Detailed methodology for immunoblotting is provided in the Supplemental Materials and Data.

### RNA isolation and quantitative PCR

All samples were homogenized with TRIzol using a PowerLyzer 24 (Qiagen), and RNA was isolated using Direct-zol RNA MiniPrep kit (Zymo Research, Cat. No. R2052) following the manufacturer’s directions. cDNA was generated from 500ng of RNA using the LunaScript RT Supermix kit (New England Biolabs, Cat. No. E3010L) according to the manufacturer’s directions. Real-time PCR (RT-PCR) was performed on a Quantstudio 3 (ThermoFisher Scientific) using either Luna Universal qPCR master mix for Sybr Green primers (New England Biolabs, Cat. No. M3003X) or Taqman Fast Advanced Master mix (ThermoFisher Scientific, Cat. No. 4444557). All primers and Taqman probes used in this work are listed in Supplemental Table 1 and the Major Resources Table. Relative gene expression was calculated using 2^-ΔΔCT^ from the relevant control group.

### Muscle cell culture

Murine C2C12 myoblasts were obtained from ATCC (Cat. No. CRL-1772) myotubes were treated with either dimethyl sulfoxide (DMSO) or uremic metabolites that are known AHR ligands (L-kynurenine, kynurenic acid, or indoxyl sulfate) for 18 hours. To explore the angiogenic secretome in C2C12 myotubes treated with DMSO or indoxyl sulfate, 25μl of undiluted conditioned media was analyzed using a custom Milliplex magnetic bead panel (Millipore-Sigma) using Luminex Magpix® multiplex analyzer and quantified using Belysa Immunoassay Curve Fitting Software (Millipore-Sigma). Primary human muscle progenitor cells (human myoblasts) were derived from fresh muscle biopsy samples as previously described ^39^ and treated with either dimethyl sulfoxide (DMSO) or uremic metabolites that are known AHR ligands (L-kynurenine, kynurenic acid, or indoxyl sulfate) for 18 hours.

### snRNA sequencing

Nuclei were isolated from the tibialis anterior muscle of male AHR^fl/fl^ and AHR^mKO^ (n=3/group) with CKD five days after FAL surgery using the Chromium nuclei isolation kit (10x Genomics, Cat. No. PN-100494). snRNA-seq libraries were generated using Chromium Next GEM Single Cell 3’ HT Reagent kits v3.1 (10x Genomics, Cat. No. PN-100370). Pooled libraries were sequenced on an Illumina NovaSeq 6000 with a S4 flow cell and 2 x 150bp pair-end reads (Azenta Life Sciences).

### snRNA-seq data processing and analysis

A detailed description of data processing and analysis is provided in the Supplementary Materials and Data. All snRNAseq data have been deposited in the Gene Expression Omnibus (https://www.ncbi.nlm.nih.gov/geo/) under accession number GSE217751.

### RNA sequencing

RNA sequencing was performed on total RNA extracted from the gastrocnemius muscle of male mice treated with AAV9-HSA-GFP and AAV9-HSA-CAAHR. A detailed description of data processing and analysis is provided in the Supplementary Materials and Data. The raw bulk RNA sequencing data have been deposited in NCBI’s Gene Expression Omnibus and can be accessed using accession number GSE225607.

### Statistical analysis

All data are presented as the mean ± standard deviation (SD). Normality of data was tested with the Shapiro-Wilk test and inspection of QQ plots. Data involving comparisons of two groups were analyzed using a student’s *t*-test when sample sizes were greater than six and normally distributed. Data involving comparisons of two groups with sample sizes less than six were analyzed using non-parametric Mann-Whitney tests. Data involving comparisons of more than two groups were analyzed using either one-way ANOVA or two-way ANOVA with Sidak’s post hoc testing for multiple comparisons when significant interactions were detected. Mixed-effects analysis was used to determine the impact of sex, diet, time, and genotype on gastrocnemius and paw perfusion recovery. In all cases, *P* < 0.05 was considered statistically significant. All statistical testing was conducted using GraphPad Prism software (version 9.0).

## RESULTS

### CKD and Uremic Toxin Exposure Result in Significant AHR Activation in Human and Mouse Skeletal Muscle

To explore whether the uremic milieu caused by CKD resulted in activation of the AHR pathway in skeletal muscle, we obtained muscle biopsy specimens from the gastrocnemius of non-PAD adult controls, PAD patients without CKD, and PAD patients with CKD. Quantitative PCR analysis demonstrated significantly higher mRNA expression of classical AHR-dependent genes (*Cyp1a1*, *Cyp1b1*, and *Aldh3a1*) in PAD patients with CKD when compared to either PAD patients with normal renal function (*P*<0.03 for all three genes) or non-PAD adult controls (*P*<0.01 for all three genes) (**Figure 1A**). Clinical and physical characteristics of these patients are shown in Table 1. Consistent with these clinical data, gastrocnemius skeletal muscle from C57BL6J mice fed an adenine-supplemented diet to induce CKD and subjected to femoral artery ligation (FAL) displayed significantly increased mRNA expression of *Ahr* and *Cyp1a1* (**Figure 1B**). Serum metabolomics of adenine-fed CKD mice confirmed a significant increase in tryptophan-derived uremic metabolites, including indoxyl sulfate, L-kynurenine, and kynurenic acid, that are known AHR ligands (**Figure 1C**). Given that the muscle specimens from human and mice contain other non-muscle cell types, we next explored the potential for AHR activation in isolated muscle cell culture systems. Acute (18h) exposure of murine (C2C12) myotubes or human primary myotubes to indoxyl sulfate, L-kynurenine, and kynurenic acid resulted in significant increases in *Cyp1a1* mRNA levels (**Figure 1D**), a classical cytochrome P450 enzyme that is transcriptionally regulated by the AHR. Taken together, these data provide conclusive evidence of AHR activation in skeletal muscle exposed to the CKD condition.

**Figure 1.**
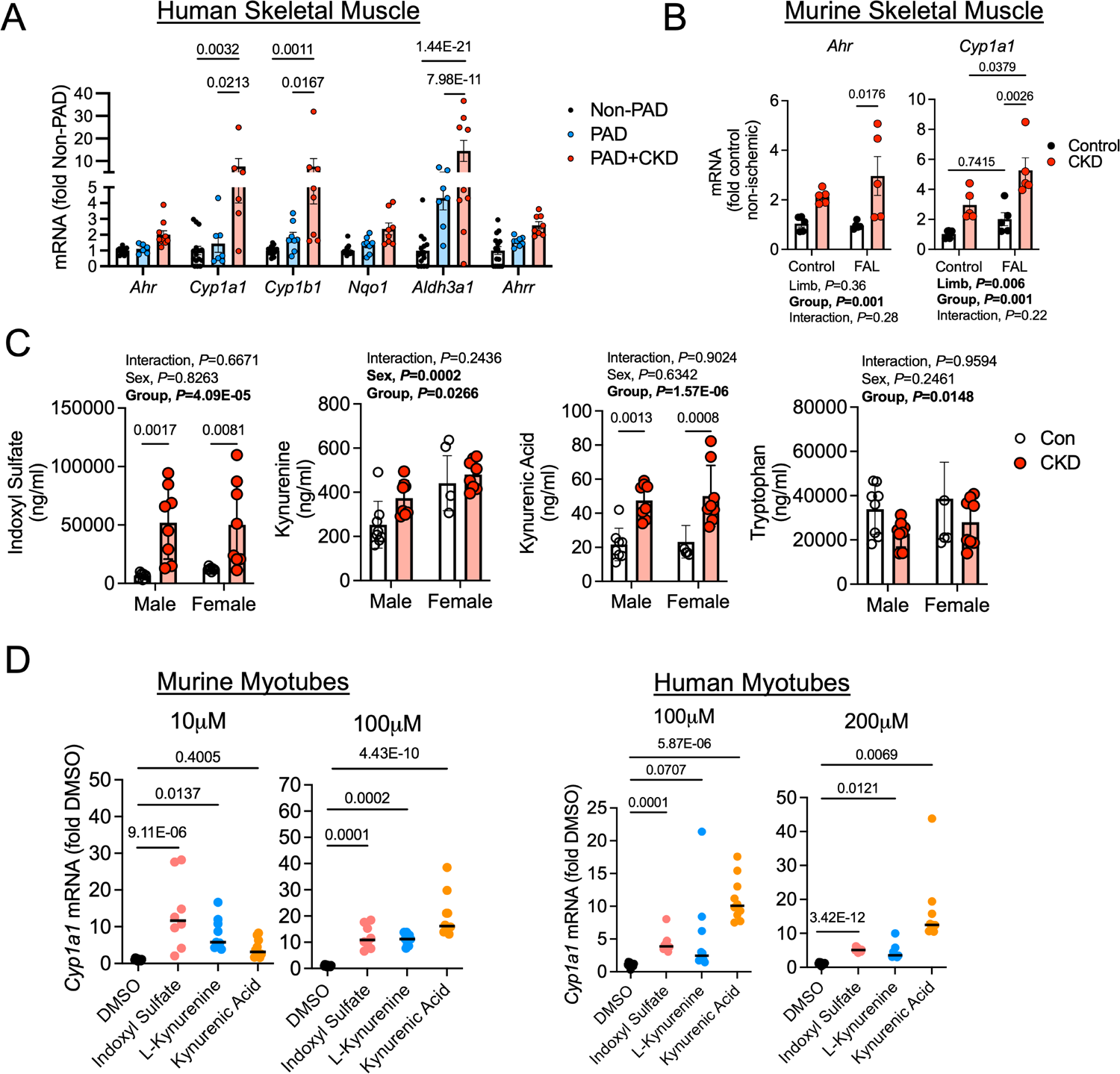
Evidence of AHR activation by uremic metabolites in human and mouse skeletal muscle. (**A**) Relative mRNA levels of genes within the AHR signaling pathway in gastrocnemius muscle specimens from Non-PAD adults (n=15) and PAD patients with (n=10) and without CKD (n=8). (**B**) Relative expression of *Ahr* and *Cyp1a1* in the gastrocnemius muscle of mice with and without CKD in both control and ischemic (FAL, femoral artery ligation) limbs (n=5/group). (**C**) Quantified levels of tryptophan-derived AHR ligands in the serum of mice with and without CKD (n=8 male Con, 4 female Con, 8 male CKD, 8 female CKD). (**D**) qPCR analysis of relative mRNA levels of *Cyp1a1*, which is transcriptionally regulated by the AHR, following acute treatment with tryptophan-derived AHR ligands shown to accumulate in CKD in both murine (n= 10 DMSO, 8 indoxyl sulfate, 9 L-kynurenine, and 10 kynurenic acid) and human cultured myotubes (n=10 DMSO, 9 indoxyl sulfate, 10 L-kynurenine, and 10 kynurenic acid). Panels A and D were analyzed using a one-way ANOVA with Sidak’s post hoc testing. Panels A and D were analyzed using a two-way ANOVA with Sidak’s post hoc testing. \**P*<0.05, \*\**P*<0.01, \*\*\**P*<0.001, \*\*\*\**P*<0.0001. Error bars represent the standard deviation.

**Table 1:** Physical and Clinical Characteristics of Human Patients.

| Characteristic | Non-PAD<br>(N=15) | PAD<br>(N=8) | PAD+CKD<br>(N=10) | P value<br>(X<sup>2</sup> or<br>ANOVA) |
| --- | --- | --- | --- | --- |
| Mean age (SD) – yr. | 70.9 (7.7) | 66.0 (6.7) | 68.4 (3.5) | 0.228<sup>A</sup> |
| Female sex – no. (%) | 6 (40) | 2 (25) | 3 (30) | **0.044** |
| Overweight/Obese (BMI ≥ 25) – no. (%) | 11 (73) | 5 (63) | 7 (70) | 0.535 |
| Ankle-brachial index (ABI) – (SD) | 1.06 (0.12) | 0.61 (0.14) | 0.51 (0.16) | **<0.001<sup>A</sup>** |
| eGFR (mL/min/1.73*m<sup>2</sup>) | >60 | 82.3 (18.6) | 27.9 (17.3) | **<0.001<sup>B</sup>** |
| Medical history – no. (%) |  |  |  |  |
| Diabetes mellitus type I or II | 6 (40) | 3 (38) | 7 (70) | 0.263 |
| Hypertension | 8 (53) | 6 (75) | 10 (100) | 0.110 |
| Hyperlipidemia | 4 (27) | 7 (88) | 8 (80) | **0.004** |
| Coronary artery disease | 0 (0) | 3 (38) | 4 (40) | 0.282 |
| Congestive Heart Failure | 0 (0) | 2 (25) | 3 (30) | 0.313 |
| Tobacco use – no. (%) |  |  |  |  |
| Former smoker | 5 (33) | 8 (100) | 10 (100) | **0.002** |
| Current smoker | 0 (0) | 0 (0) | 0 (0) | N/A |
| Medication used – no. (%) |  |  |  |  |
| Aspirin | 4 (27) | 5 (63) | 6 (60) | 0.140 |
| Statin | 4 (27) | 7 (88) | 8 (80) | **<0.004** |
| ACE inhibitor | 2 (13) | 2 (25) | 4 (40) | 0.312 |
| Cilostazol | 0 (0) | 1 (13) | 2 (20) | 0.569 |
| <sup>A</sup>ANOVA was performed. <sup>B</sup>Unpaired, two-tailed *t*-test was performed on PAD patients. $\chi^2$ analysis was performed to determine differences in population proportions. <sup>T</sup>*t*-test (two-tailed) was performed. SD, standard deviation. | | | | |

### Skeletal Muscle-Specific AHR Deletion Promotes Ischemic Muscle Perfusion Recovery in Mice with CKD

Previous work has shown that chronic AHR activation reduces angiogenesis ^21,33,34,48^ and promotes atherosclerosis ^35–37^. Additionally, chronic activation of the AHR by dioxin leads to muscle weakness and atrophy ^38,49^, symptoms commonly exhibited by PAD and CKD patients. The phenotype observed in models with chronic AHR activation is similar to the PAD patient with CKD, prompting us to hypothesize that the AHR might regulate limb pathology in this condition. To explore this, we generated an inducible skeletal muscle-specific AHR knockout mouse (AHR^mKO^, **Figure 2A**). Following administration of tamoxifen, deletion of the AHR in skeletal muscle was confirmed by immunoblotting (**Figure 2B**) and DNA recombination (**Supplemental Figure 1A,B**). Next, we randomized AHR^mKO^ mice and their wildtype (AHR^fl/fl^) littermates to either a casein control (Con) or adenine-supplement diet to induce CKD for three weeks prior to subjecting the animals to FAL as an experimental model of PAD (**Figure 2C**). CKD mice, regardless of genotype, have significantly reduced glomerular filtration rate (GFR, **Figure 2D**), elevated blood urea nitrogen (BUN, **Figure 2E**), and reduced body weights (**Figure 2F**) compared to casein fed controls, confirming a substantial level of renal dysfunction. Following FAL, CKD was found to significantly impair perfusion recovery in the paw in both male and female mice, as well as the gastrocnemius in female mice (Diet effect, *P*<0.035 in all cases; **Figure 2G,H**). Significant genotype effects were detected in both the gastrocnemius and paw of male and female mice, indicating that AHR^mKO^ mice had improvements in gastrocnemius and paw perfusion recovery (measured via laser Doppler flowmetry) (**Figure 2G,H**). To assess the effects on the muscle vasculature, we labeled endothelial cells within the tibialis anterior muscle using fluorescently labeled *Griffonia Simplicifolia* lectin I isolectin B4. Fourteen days after FAL, there was a significant effect of diet in female (*P*=0.002), but not male mice, demonstrating that CKD reduced total capillary density in female mice (**Figure 2I**).

**Figure 2.**
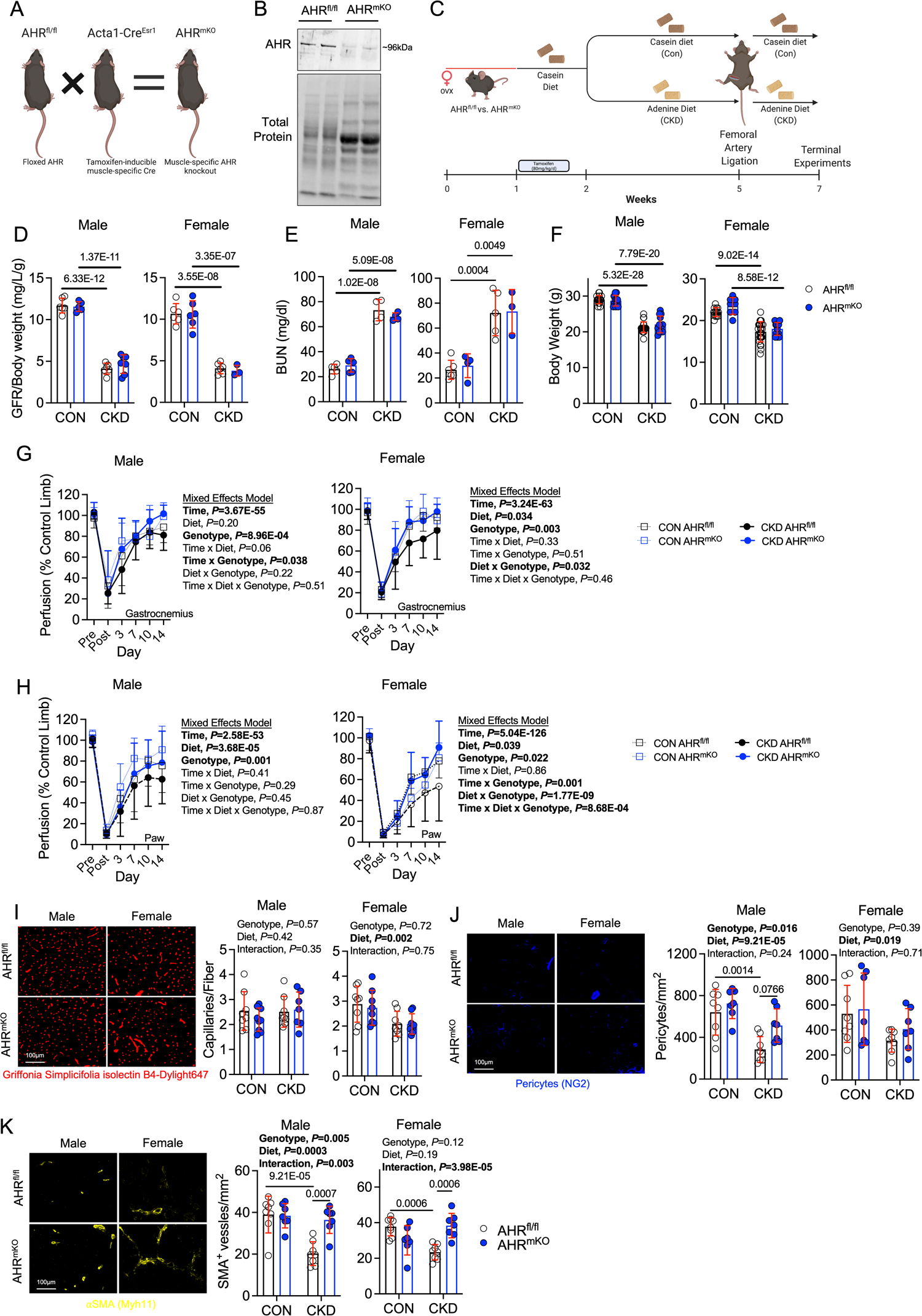
Skeletal Muscle-Specific AHR deletion Promotes Ischemic Muscle Perfusion Recovery and Arteriogenesis in Mice with CKD. (**A**) Generation of inducible, muscle-specific AHR knockout mice and (**B**) immunoblotting confirmation of AHR deletion in skeletal muscle. (**C**) Graphical depiction of the overall experiment design. (**D**) Glomerular filtration rate (GFR) relative to body weight in mice with and without CKD (n=6 AHR^fl/fl^ males/group, 5 AHR^mKO^ male CON and 7 CKD, 6 AHR^fl/fl^ female CON and 4 CKD, 6 AHR^mKO^ female CON and 3 CKD). (**E**) Quantification of blood urea nitrogen (BUN) (n=5 AHR^fl/fl^ male CON and 4 CKD, 5 AHR^mKO^ male CON and 4 CKD, 6 AHR^fl/fl^ female CON and 5 CKD, 4 AHR^mKO^ female CON and 3 CKD). (**F**) Quantification of body weights of mice with and without CKD (n=17 AHR^fl/fl^ male CON and 21 CKD, 11 AHR^mKO^ male CON and 12 CKD, 13 AHR^fl/fl^ female CON and 25 CKD, 12 AHR^mKO^ female CON and 14 CKD). (**G**) Laser Doppler flowmetry quantification of perfusion recovery in the gastrocnemius muscle (n=14 AHR^fl/fl^ male CON and 17 CKD, 13 AHR^mKO^ male CON and 13 CKD, 10 AHR^fl/fl^ female CON and 18 CKD, 9 AHR^mKO^ female CON and 8 CKD). (**H**) Laser Doppler flowmetry quantification of perfusion recovery in the paw (n=14 AHR^fl/fl^ male CON and 17 CKD, 13 AHR^mKO^ male CON and 13 CKD, 10 AHR^fl/fl^ female CON and 18 CKD, 9 AHR^mKO^ female CON and 8 CKD). Perfusion recovery was analyzed using mixed model analysis. (**I**) Quantification of capillary density in the tibialis anterior muscle (n=8 AHR^fl/fl^ males/group, 9 AHR^mKO^ male CON and 7 CKD, and n=8/genotype/diet in females). (**J**) Quantification of pericyte density in the tibialis anterior muscle (n=8 AHR^fl/fl^ males/group, 7 AHR^mKO^ males/group, and n=8 AHR^fl/fl^ ^fe^males/group, 7 AHR^mKO^ females/group). (**K**) Quantification of arteriole density in the tibialis anterior muscle (n=8 AHR^fl/fl^ males/group, 7 AHR^mKO^ males/group, and n=8 AHR^fl/fl^ ^fe^males/group, 7 AHR^mKO^ females/group). Statistical analyses performed using two-way ANOVA with Sidak’s post hoc testing for multiple comparisons when significant interactions were detected. Error bars represent the standard deviation.

Regardless of diet, there were no differences in total capillary density between AHR^mKO^ and AHR^fl/fl^ mice (**Figure 2I**). Analysis of pericyte abundance revealed a significant effect of diet in both male (*P*=0.0003) and female (*P*=0.019), demonstrating that CKD reduced pericyte abundance in the ischemic muscle (**Figure 2J**). A significant genotype effect was detected in male mice indicating the AHR^mKO^ mice had greater pericyte abundance when compared to AHR^fl/fl^ mice (*P*=0.016). Next, we examined the impact of CKD and muscle-specific AHR deletion on the abundance of arterioles by labeling muscle sections with alpha smooth muscle actin (αSMA). CKD was found to have a negative impact on arterioles abundance in both male and female AHR^fl/fl^ mice (**Figure 2K**). Strikingly, AHR^mKO^ mice with CKD, regardless of sex, display higher levels of arterioles in their ischemic limb (*P*=0.0007 and *P*=0.0006 for male and female respectively) compared to AHR^fl/fl^ mice with CKD (**Figure 2K**). In the non-ischemic control limb, there were no statistically significant adverse effects on capillary, arteriole, or pericyte density due to deletion of the AHR or the presence of CKD (**Supplemental Figure 2**).

### Skeletal Muscle-Specific Deletion of the AHR Preserves Muscle Mass, Contractile Function, and Mitochondrial Bioenergetics in Mice with CKD

Because a progressive skeletal myopathy in CKD and PAD alone is associated with adverse long-term health outcomes in patients, we next examined the impact of AHR deletion on skeletal muscle health in mice with PAD and CKD. Consistent with a wealth of preclinical and clinical literature, mice with CKD displays markedly lower muscle weights in both the ischemic limb (**Figure 3A**) and non-ischemic limb (**Supplemental Figure 3A**) compared to their casein fed counterparts with normal kidney function. Despite the catabolic state often characterized in the CKD condition, the AHR^mKO^ male mice had significantly larger masses of the ischemic tibialis anterior and extensor digitorum longus (EDL) muscles compared to AHR^fl/fl^ male mice with CKD (**Figure 3A**), although muscle masses were still lower compared to casein fed mice (*P*<0.0001). Similar trends for increased muscle masses in AHR^mKO^ mice were observed in the non-ischemic control limb muscles (**Supplemental Figure 3A).** In contrast to male mice, female AHR^mKO^ mice with CKD did not exhibit increased muscle masses in the ischemic limb when compared to female AHR^fl/fl^ mice with CKD (**Figure 3A**). Assessments of muscle force production revealed a significant diet effect in male mice (*P*=0.009), whereas the diet effect in female mice was trending but non-significant (*P*=0.08) (**Figure 3B,C**). Importantly, AHR^mKO^ male mice had significantly higher levels of absolute force at both submaximal and maximal contractile states when compared to AHR^fl/fl^ male mice with CKD (**Figure 3B**). Peak specific force levels (force normalized to area), an index of muscle quality, displayed a significant interaction (P=0.017) and subsequent post-hoc testing revealed a trending improvement in AHR^mKO^ mice with CKD compared to AHR^fl/fl^ mice with CKD (*P*=0.0573). In contrast to male mice, there were no detectable effects of genotype on muscle absolute or specific force levels (**Figure 3C**), indicating a sex-dependent effect of AHR deletion on muscle contractile function. Representative hematoxylin and eosin-stained and laminin-stained tibialis anterior muscles indicate a more preserved muscle architecture in AHR^mKO^ mice with CKD (**Figure 3D**).

**Figure 3.**
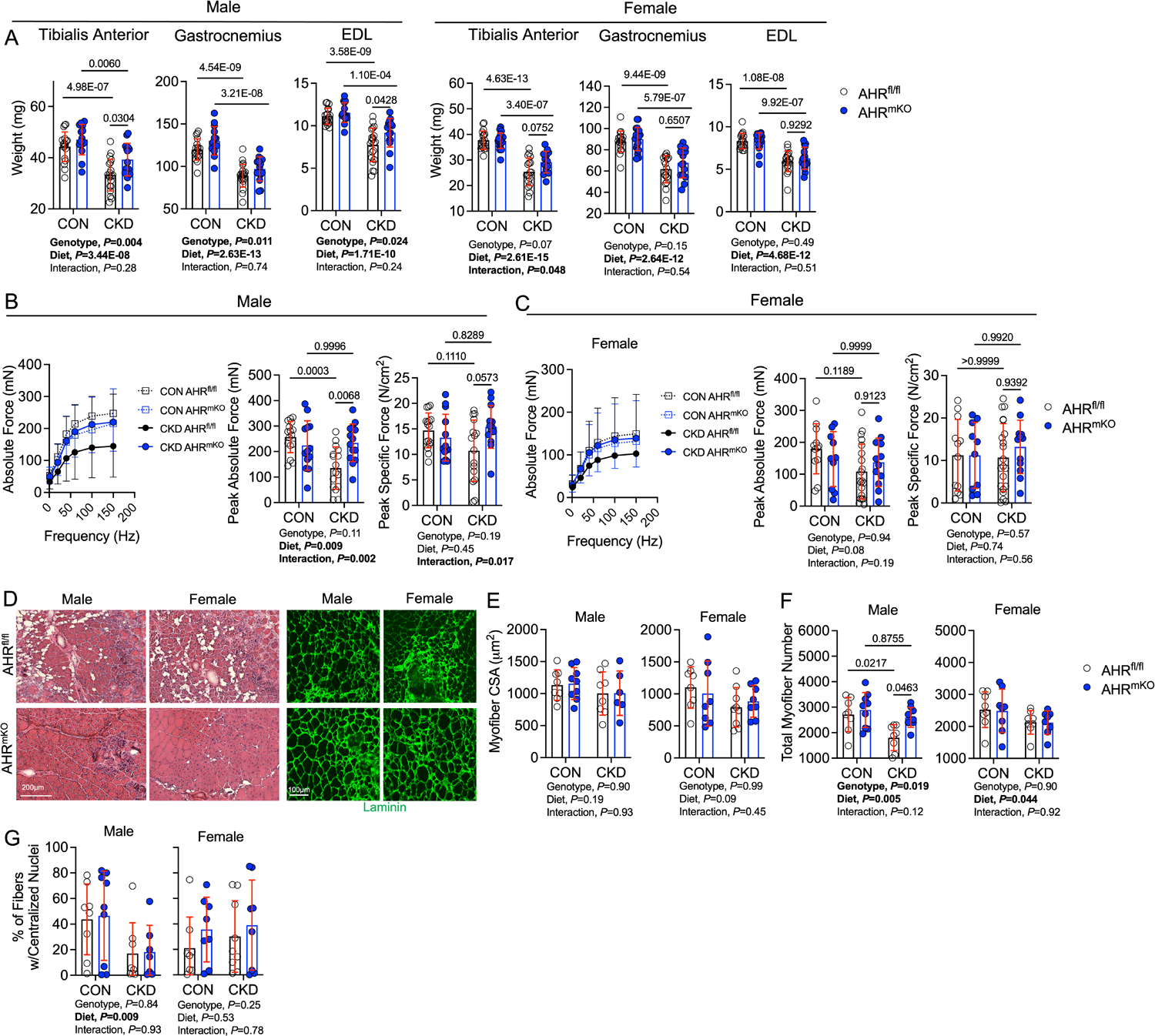
Skeletal Muscle-Specific deletion of the AHR Preserves Ischemic Muscle Mass and Contractile Function in Mice with CKD. (**A**) Quantification of muscle weights in wildtype (AHR^fl/fl^) and AHR^mKO^ mice (n=18 AHR^fl/fl^ male CON and 20 CKD, 14 AHR^mKO^ male CON and 15 CKD, 18 AHR^fl/fl^ females/group, 20 AHR^mKO^ female CON and 13 CKD). (**B**) Muscle contractile function quantification generated using *ex vivo* functional analyses of the ischemic extensor digitorum muscle in male mice (n=16 AHR^fl/fl^ male CON and 16 CKD, 13 AHR^mKO^ male CON and 14 CKD). (**C**) Muscle contractile function quantification generated using *ex vivo* functional analyses of the ischemic extensor digitorum muscle in female mice (n=12 AHR^fl/fl^ female CON and 21 CKD, 10 AHR^mKO^ female CON and 12 CKD). Specific force was calculated as the absolute force normalized to the cross-sectional area of the muscle. (**D**) Representative hematoxylin & eosin staining and laminin staining immunofluorescence from mice with CKD. (**E**) Quantification of the mean myofiber cross-sectional area of the tibialis anterior muscle (n=8 AHR^fl/fl^ males/group, 9 AHR^mKO^ male CON and 6 CKD, 8 AHR^fl/fl^ female CON and 9 CKD, 8 AHR^mKO^ females/group). (**F**) Quantification of the total fiber numbers within the tibialis anterior muscle (n=8 AHR^fl/fl^ males/group, 9 AHR^mKO^ males/group, 8 AHR^fl/fl^ females/group, 8 AHR^mKO^ females/group). (**G**) Quantification of the percentage of myofibers containing centralized nuclei within the tibialis anterior muscle (n=8 AHR^fl/fl^ males/group, 9 AHR^mKO^ male CON and 7 CKD, 8 AHR^fl/fl^ female CON and 9 CKD, 8 AHR^mKO^ females/group). Statistical analyses performed using two-way ANOVA with Sidak’s post hoc testing for multiple comparisons when significant interactions were detected. Error bars represent the standard deviation.

Interestingly, the median myofiber cross-sectional area (CSA) was not different between CKD mice with or without the AHR in the ischemic limb muscle (**Figure 3E**). However, in the non-ischemic control limb, CKD significantly decreased myofiber CSA in AHR^fl/fl^ mice but not AHR^mKO^ mice (**Supplemental Figure 3B**). Analysis of the total fiber number within the ischemic tibialis anterior muscle uncovered a significant decrease in total fibers in CKD mice irrespective of sex (**Figure 3F**). Accordant with the improved muscle mass and absolute force levels, AHR^mKO^ male, but not female, mice with CKD had significantly more myofibers compared to AHR^fl/fl^ mice with CKD (**Figure 3F**). Male mice displayed a significant diet effect for the percentage of myofibers with centralized nuclei in the ischemic limb muscle, but no effects of AHR deletion were observed in either sex (**Figure 3G**). Additionally, analysis of the fiber type distribution within the non-ischemic control limb muscle indicated that neither CKD nor deletion of AHR altered the distribution of myosin fiber types (**Supplemental Figure 3C**). Moreover, skeletal muscle-specific AHR deletion had no effect on muscle size or contractile function in non-CKD mice that received a sham surgery (**Supplemental Figure 4A-D**).

Indoxyl sulfate, one of the most well characterized uremic metabolite and AHR ligand, has been shown to impair mitochondrial respiratory function and increase reactive oxygen species (ROS) levels in mice ^22,23,32^. Coincidently, mitochondrial function has been linked to PAD limb pathobiology ^50–55^ and was identified as a potential site for the coalescence of PAD and CKD ^41^. In light of these observations, we next asked whether the AHR may have a role in mediating mitochondrial dysfunction in mice with CKD and PAD. To assess mitochondrial function, mitochondria were isolated from the ischemic gastrocnemius muscles and respirometry was performed using a creatine kinase clamp system that measures mitochondrial respiratory function across a range of energy demands (akin to a stress test) ^56–58^. When mitochondria were fueled by pyruvate and malate, AHR^mKO^ male mice with CKD displayed higher rates of oxygen consumption at all levels of energy demand assessed when compared to AHR^fl/fl^ male mice with CKD (**Figure 4A**). Quantification of conductance through the oxidative phosphorylation (OXPHOS) system (slope of oxygen consumption vs. ΔG_ATP_) uncovered a significant interaction (*P*=0.003) and post-hoc testing revealed significantly greater OXPHOS conductance in male AHR^mKO^ mice compared to male AHR^fl/fl^ mice (**Figure 4A**). Similarly, maximal uncoupled respiration (following the addition of FCCP) fueled by pyruvate/malate was significantly higher in male AHR^mKO^ mice with CKD compared to AHR^fl/fl^ male mice with CKD (*P*=0.0068, **Figure 4A**). Parallel experiments were also performed when fueling mitochondria with a medium chain fatty acid (octanoylcarnitine). These experiments revealed significant genotype effects, but no interactions between diet and genotype, indicating the AHR^mKO^ male mice had improved mitochondrial fatty acid oxidation compared to AHR^fl/fl^ male mice (**Figure 4B**). Interestingly, the effects of muscle-specific AHR deletion on mitochondrial function was absent in female mice, regardless of the kidney function or fuel source utilized (**Figure 4C,D**). Similar to contractile function, skeletal muscle-specific AHR deletion had no effect on mitochondrial function in non-CKD mice that received a sham surgery (**Supplemental Figure 4E,F**).

**Figure 4.**
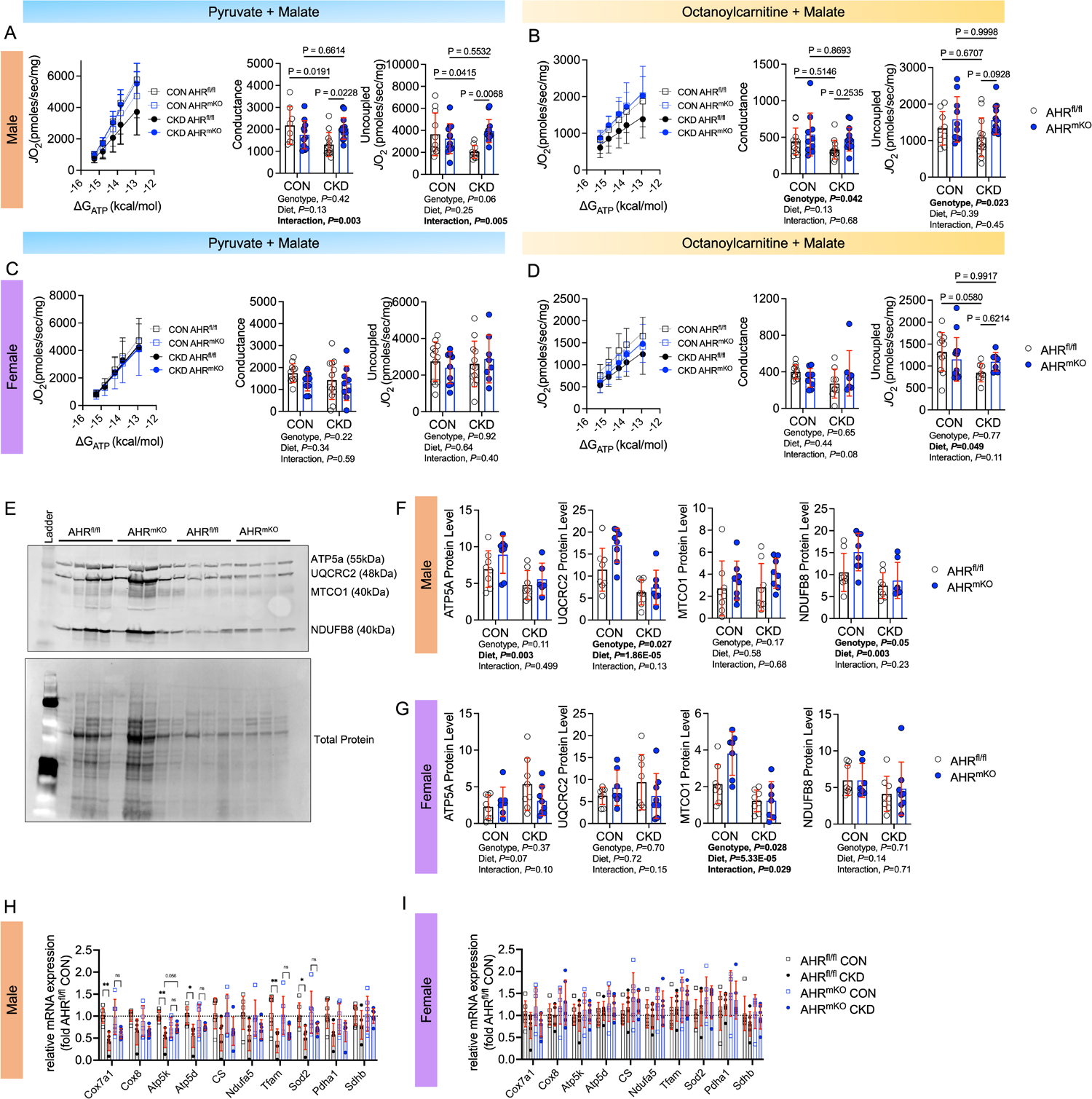
Skeletal Muscle-Specific deletion of the AHR Preserves Ischemic Muscle Mitochondrial Energetics in Mice with CKD. Muscle mitochondrial function was assessed using a creatine kinase clamp system to measure oxygen consumption (*J*O_2_) at physiologically relevant energy demands (ΔG_ATP_). (**A**) Relationship between *J*O_2_ and ΔG_ATP_ when mitochondria were fueled with pyruvate and malate and quantification of the conductance (slope of *J*O_2_ and ΔG_ATP_ relationship) and uncoupled respiration rates in male mice (n=8 AHR^fl/fl^ male CON and 12 CKD, 11 AHR^mKO^ male CON and 14 CKD). (**B**) Relationship between *J*O_2_ and ΔG_ATP_ when mitochondria were fueled with octanoylcarnitine and malate and quantification of the conductance and uncoupled respiration rates in male mice (n=11 AHR^fl/fl^ male CON and 13 CKD, 10 AHR^mKO^ male CON and 13 CKD). (**C**) Relationship between *J*O_2_ and ΔG_ATP_ when mitochondria were fueled with pyruvate and malate and quantification of the conductance and uncoupled respiration rates in female mice (n=11 AHR^fl/fl^ female CON and 13 CKD, 10 AHR^mKO^ female CON and 9 CKD). (**D**) Relationship between *J*O_2_ and ΔG_ATP_ when mitochondria were fueled with octanoylcarnitine and malate and quantification of the conductance and uncoupled respiration rates in female mice (n=11 AHR^fl/fl^ female CON and 9 CKD, 9 AHR^mKO^ female CON and 7 CKD). (**E**) Western blotting membrane detecting the abundance of mitochondrial electron transport system proteins (n=8/group/sex/genotype). (**F**) Quantification of mitochondrial protein abundance in male mice n=8/group/sex/genotype). (**G**) Quantification of mitochondrial protein abundance in female mice n=8/group/sex/genotype). (**H**) qPCR analysis of mitochondrial transcripts in male mice (n=6/group). (**I**) qPCR analysis of mitochondrial transcripts in female mice (n=6/group/sex/genotype). Statistical analyses performed using two-way ANOVA with Sidak’s post hoc testing for multiple comparisons when significant interactions were detected. Error bars represent the standard deviation.

Next, mitochondrial content/density was explored using immunoblotting for select proteins in the electron transport system (**Figure 4E**). In male mice, a significant diet effect was found for ATP5A, UQCRC2, and NDUFB8 proteins indicating that CKD reduced the abundance of these proteins (**Figure 4F,** all blot images shown in **Supplemental Figure 5**). Further to this, AHR^mKO^ mice were found to have significantly higher levels UQCRC2 and NDUFB8 protein levels (*P*=0.027 and *P*=0.05 respectively). In female mice, only MTCO1 protein levels were reduced in CKD mice (**Figure 4G**). Congruent with the findings from respirometry experiments, mitochondrial protein abundance was largely unaffected by the deletion of the AHR in female mice. At the level of the mRNA, the expression of several mitochondrial genes (*Cox7a1, Atp5k, Atp5d, Tfam, Sod2*) were significantly lower in male AHR^fl/fl^ mice with CKD compared to AHR^fl/fl^ mice without CKD (**Figure 4H**); however, these changes were attenuated in AHR^mKO^ male mice with CKD. In contrast, mRNA levels for the examined mitochondrial genes were minimally impacted by either CKD or the deletion of the AHR in the ischemic muscle of female mice (**Figure 4I**). In the non-ischemic control limb, mitochondrial gene expression was minimally affected except for *Ndufa5* and *Tfam* which were reduced in AHR^fl/fl^ male mice with CKD but significantly greater in AHR^mKO^ male mice with CKD (**Supplemental Figure 6**).

### Single Nuclei RNA Sequencing (snRNA-seq) Reveals Differences in Cell Populations and Transcriptional Profile in Ischemic Muscle of AHR^mKO^ mice

To compare different nuclei populations in AHR^fl/fl^ and AHR^mKO^ skeletal muscles from CKD/PAD mice, we performed snRNA-seq on nuclei isolated from the tibialis anterior muscle of AHR^fl/fl^ and AHR^mKO^ harvested five days after FAL surgery (**Figure 5A**). Importantly, these muscle specimens were harvested at a timepoint when limb perfusion was not different between groups (**Figure 5B**). Nuclei were isolated using gentle homogenization, purified using the Chromium nuclei isolation kit (10x Genomics), and quality determined using fluorescence microscopy. From AHR^fl/fl^ muscle, we captured 18,833 nuclei, and from AHR^mKO^ muscle we captured 12,597 nuclei with high integrity. A median of 1,853 genes per nuclei were sequenced from AHR^fl/fl^ muscle, whereas a median of 1,110 genes per nuclei were sequenced from AHR^mKO^ muscle. The total number of genes detected was 25,964 and 25,009 in AHR^fl/fl^ and AHR^mKO^ mice respectively.

**Figure 5.**
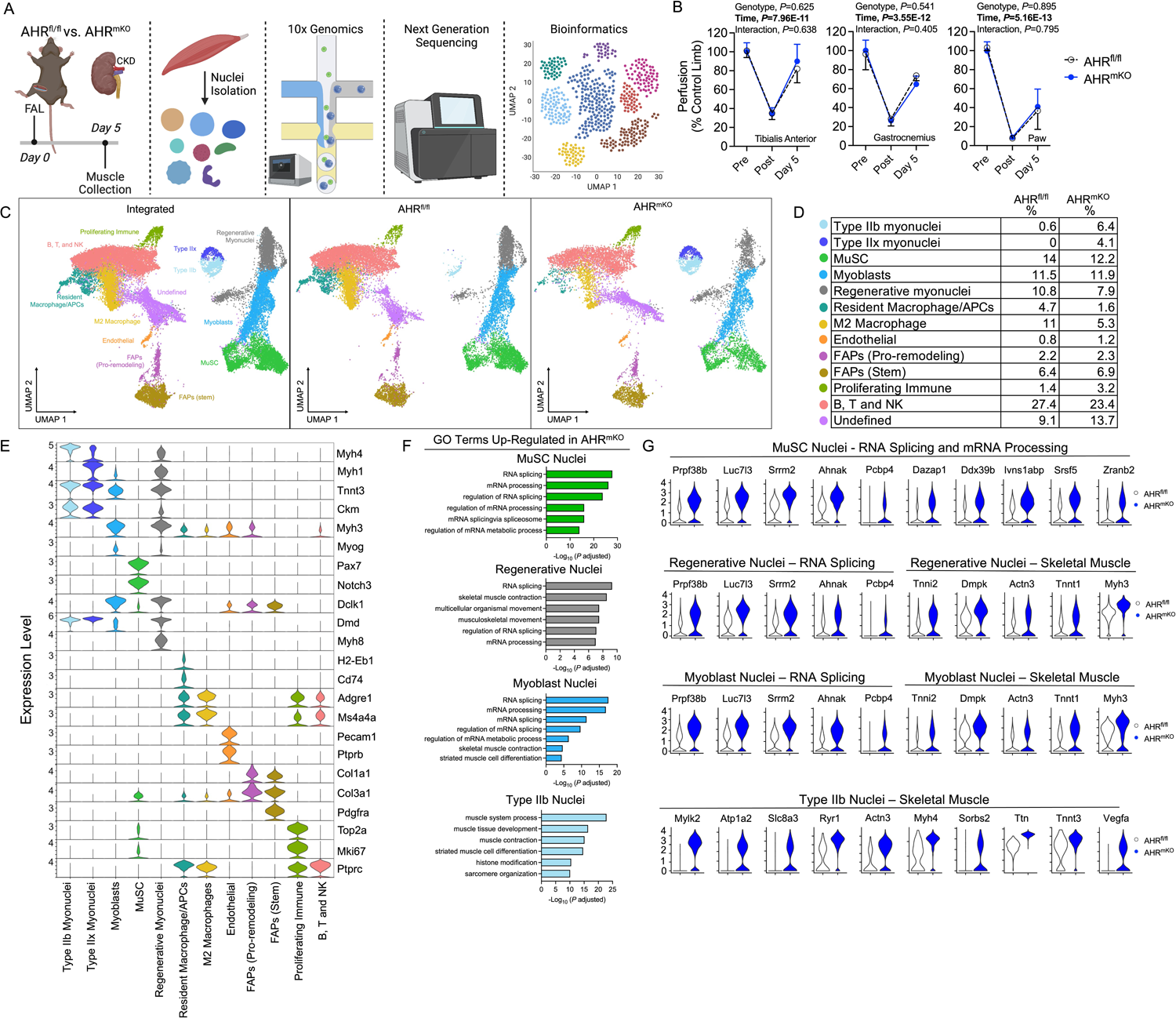
snRNA sequencing on ischemic muscle from AHR^fl/fl^ and AHR^mKO^ mice with CKD. (**A**) Schematic of the experimental design for snRNA-seq experiment. (**B**) Laser Doppler Flowmetry quantification of limb perfusion demonstrates an equivalent ischemic burden between groups. (**C**) UMAPs visualization of the integrated datasets to identify clusters, as well as UMAPs for AHR^fl/fl^ and AHR^mKO^ muscle nuclei. (**D**) Percentage of nuclei within each cluster determined by group. (**E**) Violin plots showing z-score transformed expression of selected marker genes used to determine cluster identity. (**F**) Gene ontology (GO) analysis of significantly upregulated genes in AHR^mKO^ myonuclei populations. (**G**) Violin plots showing normalized expression levels of select differentially expressed genes from top GO terms.

To compare AHR^fl/fl^ and AHR^mKO^ samples, we integrated the datasets resulting in a total of 31,430 nuclei for bioinformatic analysis including using uniform manifold approximation and projection (UMAP) to visualize and resolve different nuclear populations (**Figure 5C**). We identified 13 clusters of nuclei based upon their transcriptional profiles and assigned identities by examining normalized expression of values of top markers and known marker genes for each cluster (**Figure 5C**) and the relative percentages of each nuclei population are shown in **Figure 5D**. Violin plots for selected marker genes in each known nuclei population are depicted in **Figure 5E**. These analyses identified differences in the abundance of mature Type IIb and Type IIx myonuclei, with AHR^mKO^ having more mature myonuclei compared to AHR^fl/fl^ (**Figure 5C,D**). Considering the time course of muscle regeneration relative to the ischemic injury (day 5 post-FAL), this finding suggests that deletion of the AHR in skeletal muscle promotes the survival of mature myonuclei during severe limb ischemia in mice with CKD. Next, we explored differentially expressed genes and gene ontology (GO) pathways in myonuclei populations to uncover potential differences in transcriptional activity in AHR^mKO^ mice. We identified that pathways related to RNA splicing, mRNA processing, and mRNA metabolic process, which are involved in muscle growth processes, were upregulated in AHR^mKO^ muscle stem cell (MuSC), Regenerative, and Type IIx myonuclei populations (**Figure 5F**). In mature Type IIb and Type IIx myonuclei, upregulated pathways in AHR^mKO^ mice included ‘*muscle contraction’* and ‘*generation of precursor metabolites and energy*’, both of which are consistent with the observed preservation of muscle force production and mitochondrial function in AHR^mKO^ mice with CKD following FAL surgery. Violin plots with expression levels of select differentially expressed genes involved in the top GO terms are shown in **Figure 5G**.

### Skeletal Muscle-Specific Deletion of the AHR Preserves Paracrine Signaling Between the Muscle and Vasculature in Mice with CKD

To investigate the potential intercellular communications that could explain the improved perfusion recovery in AHR^mKO^ mice with CKD (**Figure 2G,H**), ligand-receptor interactions were calculated from snRNA sequencing data using CellChat ^59^. Circle plots showing the overall intercellular communication occurring in AHR^fl/fl^ and AHR^mKO^ muscles are shown in **Figure 6A**. Circle sizes represent the number of cells, whereas the edge width represents the communication probability and edge color is set according to cell type. For the inferred angiogenic pathways, *Vegf* and *Notch* signaling pathways were present in AHR^mKO^ muscles but were absent in AHR^fl/fl^ muscles (**Figure 6B**). Moreover, the VEGF pathway communication was strong between the endothelial nuclei and mature Type IIb and IIx myonuclei populations of AHR^mKO^ muscles. Circle plots for communications of each individual cell type are shown in **Supplemental Figure 7**. All significant signaling pathways were then ranked based upon their differences in relative overall information flow between AHR^fl/fl^ and AHR^mKO^ muscles (**Figure 6C**). Enriched angiogenic pathways in AHR^mKO^ muscles included *Vegf*, *Notch*, and *Pecam1*. Enriched pathways in AHR^fl/fl^ muscles included TGF-beta, collagen, granulin, and interestingly, angiopoietin. Analysis of the outgoing signaling patterns of secreting cells indicated that Type IIb and IIx myonuclei were the likely source of the angiogenic growth factor VEGF in AHR^mKO^ muscles, whereas NOTCH ligands were most likely secreted from FAPs and myoblasts (**Figure 6D**). Analysis of the incoming communication pathways indicated that secreted VEGF was most likely interacting with VEGF receptor in endothelial cells of AHR^mKO^ muscles (**Figure 6D**).

**Figure 6.**
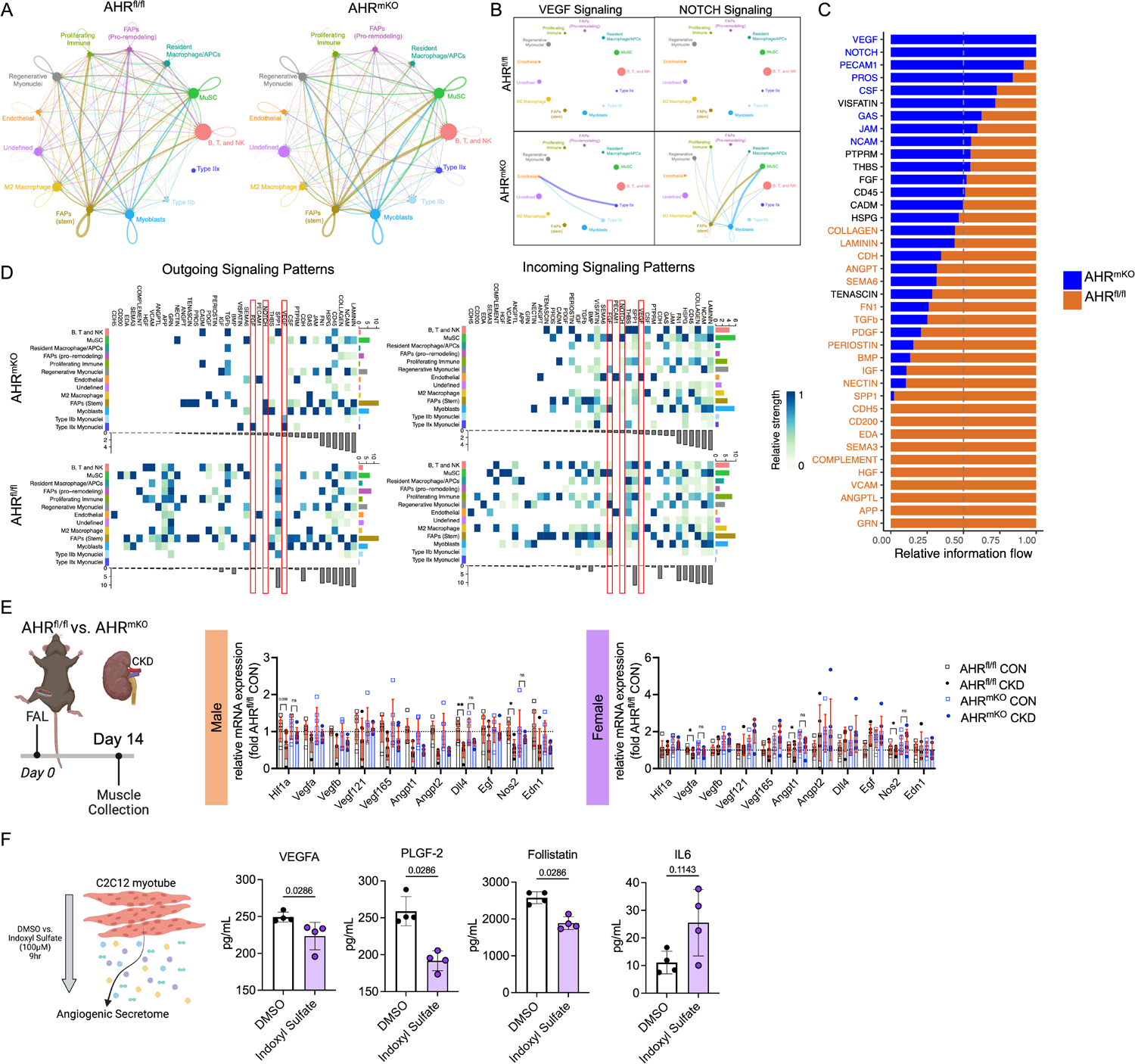
AHR^mKO^ mice with CKD have preserved paracrine vasculogenic signaling between myofibers and vascular cells. (**A**) Circle plots showing the overall intercellular communication occurring in AHR^fl/fl^ and AHR^mKO^ muscles. (**B**) Circle plots showing VEGF and NOTCH signaling communication in AHR^fl/fl^ and AHR^mKO^ muscles. (**C**) Ranked significant ligand-receptor communications for relative information flow between AHR^fl/fl^ and AHR^mKO^ muscles. (**D**) Enriched outgoing and incoming signaling patterns according to cell type based on signaling strength. (**E**) Targeted angiogenic and vascular reactivity qPCR analysis on total RNA from the ischemic hindlimb muscles (n=6/group/sex/genotype). (**F**) Analysis of the protein abundance of secreted angiogenic growth factors in conditioned media from C2C12 myotubes (n=4/group). Analyses in A-D were performed using CellChat. Panel E analyses involved a two-way ANOVA with Sidak’s post hoc testing for multiple comparisons when significant interactions were detected. Panel F was analyzed with a Mann-Whitney test. Error bars represent the standard deviation.

Using traditional bulk qPCR in RNA isolated from ischemic muscle, the Hif1-alpha expression levels tended to be lower in AHR^fl/fl^ male mice with CKD (*P*=0.056) but was not different in AHR^mKO^ mice with CKD (**Figure 6E**). Considering that AHR forms a heterodimer with the aryl hydrocarbon receptor nuclear translocator *(Arnt*), also known as Hif1-beta, this finding suggests that chronic AHR activation may disrupt hypoxia signaling cascades through interactions of the AHR with hypoxia inducible transcription factors. Several angiogenic genes displayed a non-significant elevation of expression including *Vegfa, Vegf121, Vegf165, Dll4, and Egf* in male AHR^mKO^ mice with CKD (**Figure 6E**). In female mice, disruptions in Hif1-alpha were not observed (**Figure 6E**). Female AHR^fl/fl^ mice with CKD were found to have significantly lower expression angiogenic growth factors *Vegfa* and *Angpt1*, but these effects were abolished in AHR^mKO^ females (**Figure 6E**). Similar to male mice, female AHR^mKO^ also displayed non-significant increases in several other angiogenic genes including *Vegf121* and *Vegf165*. Regarding vasomotor tone, both male and female AHR^fl/fl^ mice with CKD, but not AHR^mKO^, displayed reduced expression of *Nos2* which has a prominent role in regulating vasodilation (**Figure 6E**). Parallel analysis of these mRNAs in the non-ischemic limb demonstrated that AHR^mKO^ mice with CKD has greater expression of several angiogenic transcripts in male (*Vegf121, Angpt1, Angpt2, Egf*) and female (*Vegf121, Vegf165, Egf*) compared to their AHR^fl/fl^ correspondents with CKD (**Supplemental Figure 8**). Further to this, male AHR^mKO^ mice with CKD had higher expression levels of *Nos2* in their non-ischemic limb muscle compared to male AHR^fl/fl^ mice with CKD (**Supplemental Figure 8**). These results suggest that angiogenic and vasodilatory gene expression was likely higher in the AHR^mKO^ limb muscle prior to FAL surgery which likely contributed to the improved vascular recovery observed by laser Doppler flowmetry. To confirm a functional impact on paracrine signaling, we measured angiogenic growth factor levels in conditioned media taken from C2C12 myotubes treated with either vehicle of indoxyl sulfate to activate the AHR. Results from this experiment demonstrated that myotubes with AHR activation secreted significantly less VEGFa, placental growth factor 2 (PLGF-2), and follistatin, as well as a trending increase in IL6 (**Figure 6F**). The totality of these findings demonstrates that AHR activation in muscle diminishes paracrine angiogenic/vasculogenic signaling.

### Expression of a Constitutively Active AHR Decreases Capillary Density and alters Angiogenic Gene Expression in Male Mice with Normal Kidney Function

Because CKD results in the retention/accumulation of many uremic solutes that are not AHR ligands ^19,20^ we sought to evaluate the role of AHR activation in mice with normal renal function. To accomplish this, we generated a constitutively active AHR (CAAHR) by deletion of the ligand binding domain (amino acids 277-418) ^60^ and used adeno-associated virus (AAV) to express locally in the myofibers of the hindlimb driven by the human skeletal actin (HSA; ACTA1 gene) promoter (**Figure 7A,B**). In this experiment, two control groups were utilized that received either a green fluorescent protein (GFP) or the full coding sequence of the AHR which requires ligands for activation (AAV-HSA-AHR). qPCR analysis of muscle confirmed that both AAV-HSA-AHR and AAV-HSA-CAAHR treated mice had elevated expression of the *Ahr*, whereas *Cyp1a1* mRNA levels were only increased in the AAV-HSA-CAAHR treated muscle, confirming constitutive AHR activation (**Figure 7C**). Interestingly, a significant interaction effect was detected for *Cyp1a1* (*P*=0.001) indicating that females had greater expression than males treated with AAV-HSA-CAAHR. Following 12-weeks of AAV expression, FAL was performed and laser Doppler flowmetry assessments did not identify any significant alterations in perfusion recovery of the gastrocnemius muscle (**Figure 7D**). However, histological quantification of capillaries revealed that male mice treated with AAV-HSA-CAAHR had lower capillary density compared to GFP treated controls, whereas females treated with AAV-HSA-CAAHR had higher capillary density compared to GFP treated controls (**Figure 7E**). Quantification of pericyte abundance in the ischemic muscle revealed a similar density of pericytes in AAV-HSA-GFP and AAV-HSA-CAAHR treated mice (Figure 7F). A trending, but non-significant, decrease in arteriole abundance was observed in AAV-HSA-CAAHR mice (*P*=0.054, **Figure 7G**). RNA sequencing performed on total RNA isolated from the gastrocnemius muscle mice treated with either AAV-HSA-GFP or AAV-HSA-CAAHR showed that treatment with AAV-HSA-CAAHR decreased the expression of angiogenic growth factors (*Vegfd, Fgf2, Egr3*), genes involved in remodeling of the extracellular matrix (*Mmp19, Mmp9, Timp3*), vascular cell adhesion (*Vcam1*), and vasoreactivity (*Ptgs2*) (**Figure 7H**). Taken together, these results indicate that constitutive activation of the AHR in muscle in mice with normal renal function is sufficient to alter the angiogenic response to ischemia.

**Figure 7.**
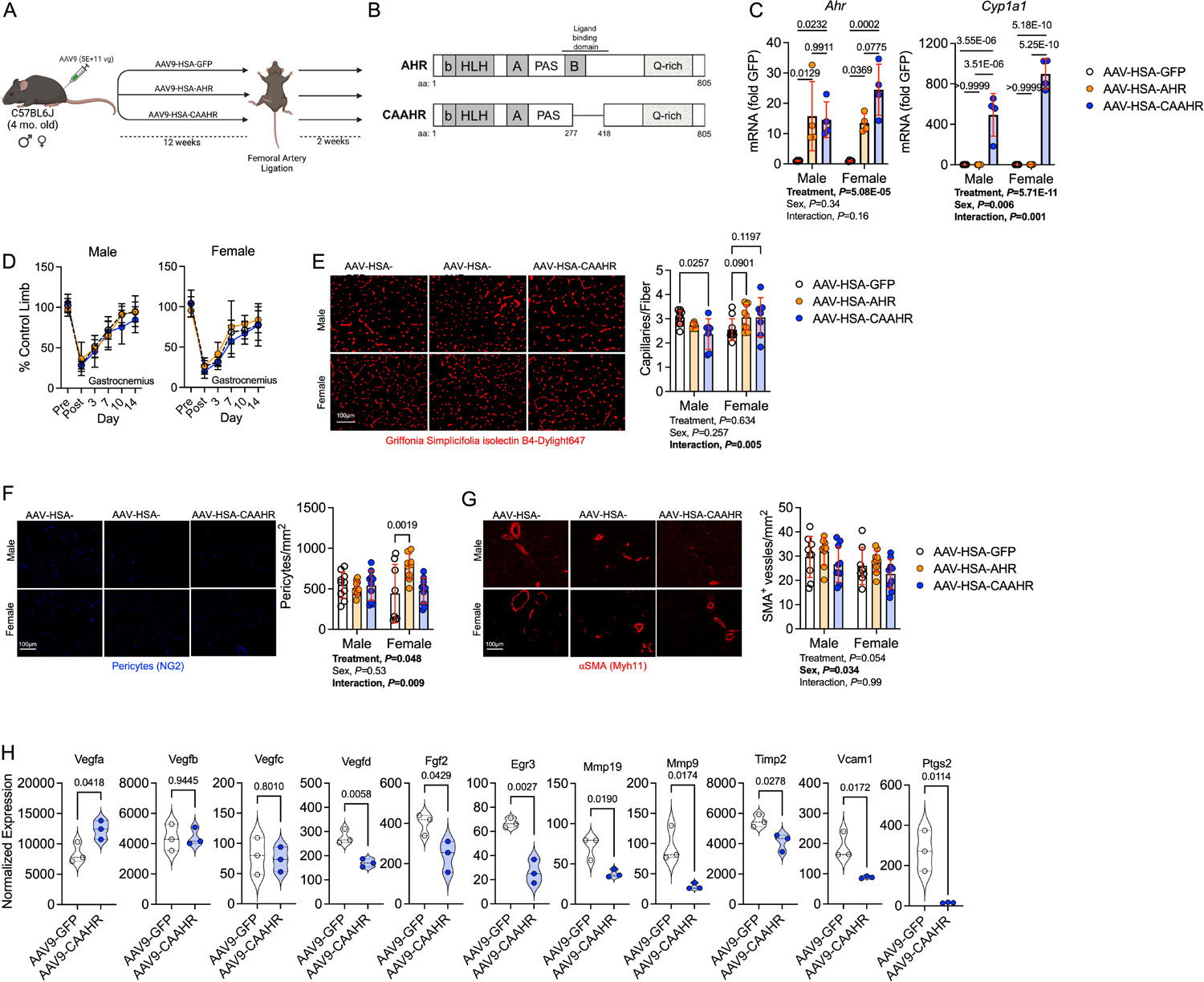
Expression of a Constitutively Active AHR Decreases Capillary Density and alters Angiogenic Signaling in Male Mice with Normal Kidney Function. (**A**) Schematic of the experimental design involving local intramuscular delivery of AAV9 to manipulate the AHR pathway in mice with normal kidney function. (**B**) Generation of a mutant AHR that displays constitutive transcriptional activation (CAAHR) via deletion of the ligand binding domain. (**C**) Quantification of mRNA levels of *Ahr* and *Cyp1a1* confirm constitutive activation in mice treated with the CAAHR driven by the human skeletal actin (HSA) promoter (n=4/group/sex). Analysis performed using one-way ANOVA with Dunnett’s post hoc testing. (**D**) Laser Doppler perfusion recovery in the gastrocnemius muscle was not different across treatment groups or sexes (n=10/group/sex). (**E**) Representative images of the tibialis anterior muscle labeled for endothelial cells and quantification of capillary density (n=10 AAV-HSA-GFP/sex, 6 male and 9 female AAV-HSA-AHR, and 7 AAV-HSA-CAAHR/sex). (**F**) Representative images of the tibialis anterior muscle labeled for perictes and quantification of pericyte density (n=10 male and 8 female AAV-HSA-GFP, 10 male and 9 female AAV-HSA-AHR, and 10 AAV-HSA-CAAHR/sex). (**G**) Representative images of the tibialis anterior muscle labeled for arterioles and quantification of arteriole density (n=9 male and 8 female AAV-HSA-GFP, 9male and 10 female AAV-HSA-AHR, and 10 AAV-HSA-CAAHR/sex). (**H**) Angiogenic and vasoreactive associated gene expression in AAV-HSA-GFP and AAV-HSA-CAAHR muscle quantified via bulk RNA sequencing analysis (n=3 male mice per group). Analysis of panels C-G was performed using two-way ANOVA with Sidak’s post hoc testing for multiple comparisons. Analysis in Panel H involved false-discovery rate corrected t-test. Error bars represent the standard deviation.

### Expression of a Constitutively Active AHR in Myofibers Exacerbates Ischemic Myopathy in Mice with Normal Kidney Function

Next, we examined whether chronic AHR activation in skeletal myofibers in the absence of renal dysfunction, was sufficient to drive myopathy in mice subjected to FAL. Consistent with the catabolic phenotype of CKD, the muscle weights of mice treated with AAV-HSA-CAAHR were significantly lower than those treated with either AAV-HSA-GFP or AAV-HSA-AHR (**Figure 8A**). Assessments of muscle contractile function demonstrated a further reduction in specific force levels, demonstrating that expression of the CAAHR severely diminishes muscle quality (force per area/size) (**Figure 8B**). Notably, the effect sizes of CAAHR expression on muscle function were larger in male mice compared to females (mean difference in peak specific force between GFP and CAAHR = 6.1 vs. 3.7 N/cm^2^ for males and females respectively). Histological analysis of ischemic tibialis anterior muscles uncovered pathological muscle remodeling in mice treated with CAAHR including increased numbers of centrally nucleated myofibers (males only), increased non-muscle areas (both sexes), and increased ischemic lesion areas (both sexes) (**Figure 8D**). Furthermore, AAV-HSA-CAAHR treated mice had a significantly smaller median myofiber cross-sectional area (CSA) compared to AAV-HSA-GFP controls subjected to FAL (**Figure 8E**).

**Figure 8.**
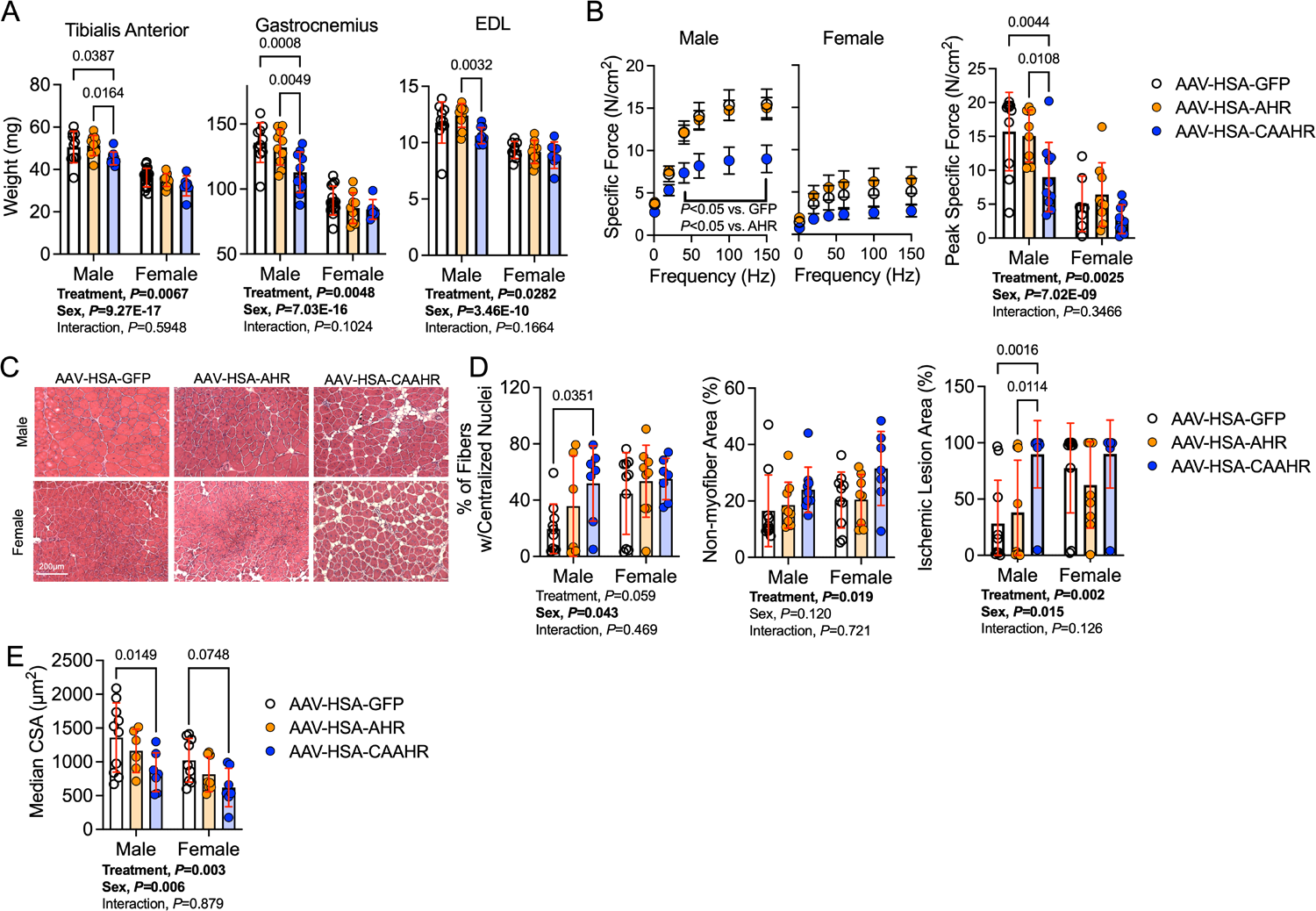
Expression of a Constitutively Active AHR in Myofibers Exacerbates Ischemic Myopathy in Mice with Normal Kidney Function. (**A**) Quantification of muscle weights from the ischemic limb of treated mice (n=10 male and 14 female AAV-HSA-GFP, 10 male and 9 female AAV-HSA-AHR, and 10 male and 7 female AAV-HSA-CAAHR). (**B**) Force frequency curves generated using *ex vivo* functional analyses of the ischemic extensor digitorum muscle in male and female mice with CKD. Specific force was calculated as the absolute force normalized to the cross-sectional area of the muscle (n=10 male and 7 female AAV-HSA-GFP, 10 male and 9 female AAV-HSA-AHR, and 10 male and 9 female AAV-HSA-CAAHR). (**C**) Representative hematoxylin & eosin staining of the ischemic tibialis anterior muscles. (**D**) Quantification of the histopathology of the ischemic tibialis anterior muscles (n=10 AAV-HSA-GFP/sex, 9 AAV-HSA-AHR/sex, and 10 AAV-HSA-CAAHR/sex). (**E**) Quantification of the mean myofiber area in the ischemic tibialis anterior muscles (n=10 male and 9 female AAV-HSA-GFP, 6 male and 8 female AAV-HSA-AHR, and 7 AAV-HSA-CAAHR/sex). Analysis performed using two-way ANOVA with Sidak’s post hoc testing for multiple comparisons. Error bars represent the standard deviation

### Expression of a Constitutively Active AHR in Myofibers Decreases OXPHOS Function in Ischemic Muscle of Mice with Normal Kidney Function

To explore the impact of CAAHR expression on ischemic muscle mitochondrial OXPHOS function, we isolated mitochondria from the ischemic gastrocnemius muscle, energized them with a mixture of carbohydrates (pyruvate) and a medium chain fatty acid (octanoylcarnitine), and assessed the respiration rate across increasing levels of energy demand (**Figure 9A**). Quantification of OXPHOS conductance unveiled a significant defect in muscle mitochondrial OXPHOS in mice treated with AAV-HSA-CAAHR when compared to either GFP- or AHR-treated controls (**Figure 9B**). Following the addition of a mitochondrial uncoupler (the protonophore, FCCP), the maximal respiratory capacity (**Figure 9C**) was also significantly lower in AAV-HSA-CAAHR treated ischemic muscle from mice with normal function. These results are consistent with the preservation of muscle mitochondrial OXPHOS function in AHR^mKO^ mice with CKD/PAD (**Figure 4**) and provides compelling evidence indicating that chronic AHR activation alters skeletal muscle mitochondrial function.

**Figure 9.**
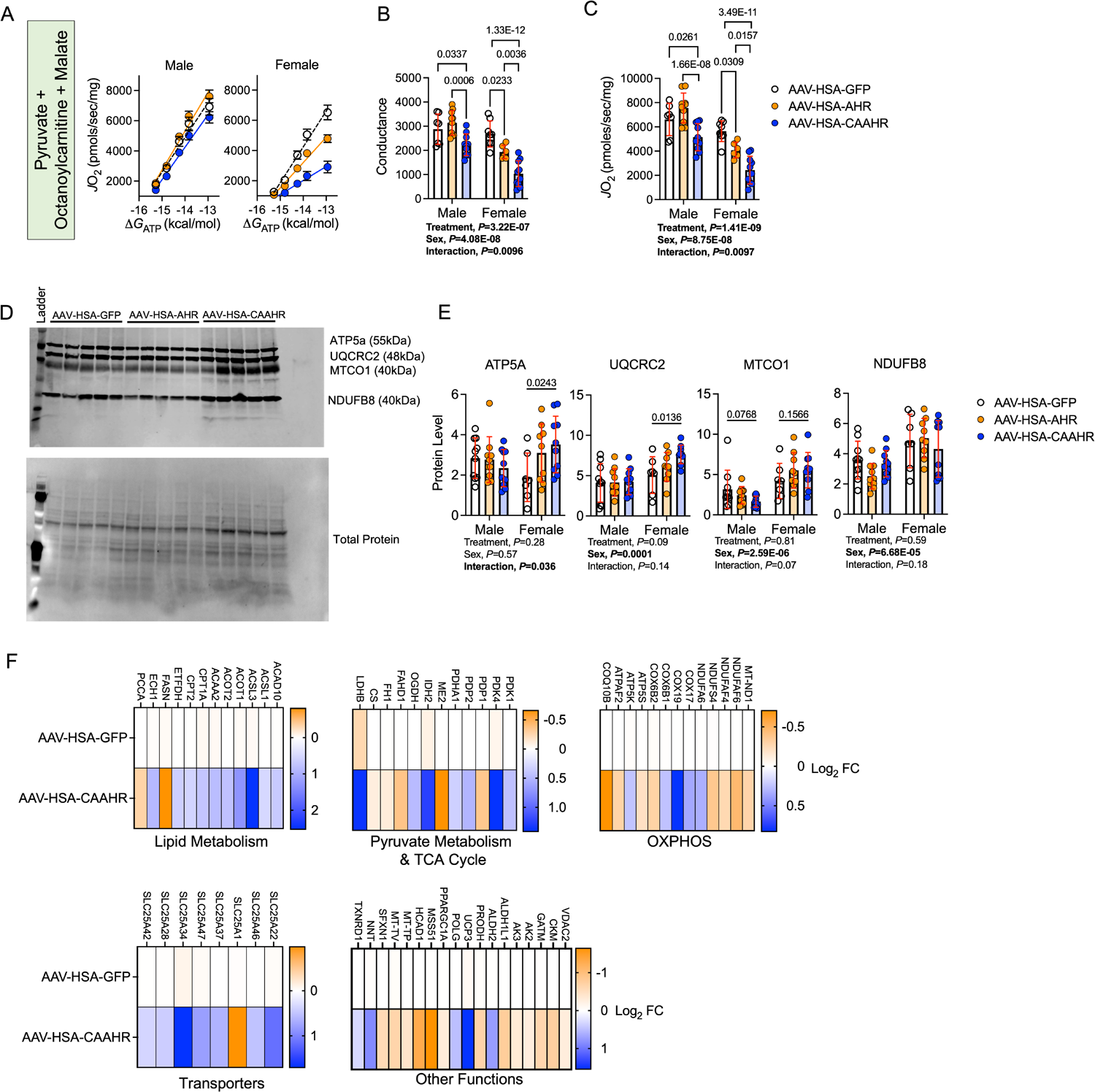
Expression of a Constitutively Active AHR in Myofibers Decreases OXPHOS Function in Ischemic Muscle of Mice with Normal Kidney Function. Muscle mitochondrial function was assessed using a creatine kinase clamp system to measure oxygen consumption (*J*O_2_) at physiologically relevant energy demands (ΔG_ATP_). (**A**) Relationship between *J*O_2_ and ΔG_ATP_ when mitochondria were fueled with pyruvate, octanoylcarnitine, and malate (n=7 male and 8 female AAV-HSA-GFP, 10 male and 6 female AAV-HSA-AHR, and 10 AAV-HSA-CAAHR/sex). (**B**) Quantification of the conductance (slope of *J*O_2_ and ΔG_ATP_ relationship) uncovered a significant treatment effect (n=7 male and 8 female AAV-HSA-GFP, 10 male and 6 female AAV-HSA-AHR, and 10 AAV-HSA-CAAHR/sex). (**C**) Maximal uncoupled respiration following the addition of a mitochondrial protonophore/uncoupler (FCCP) (n=7 male and 8 female AAV-HSA-GFP, 10 male and 6 female AAV-HSA-AHR, and 10 AAV-HSA-CAAHR/sex). (**D**) Western blotting membrane detecting the abundance of mitochondrial electron transport system proteins. (**E**) Quantification of mitochondrial protein abundance in male and female mice (n=10 male and 7 female AAV-HSA-GFP, 10 male and 9 female AAV-HSA-AHR, and 10 AAV-HSA-CAAHR/sex). (**F**) Mitochondrial associated gene expression (Log_2_ fold change) in AAV-HSA-GFP and AAV-HSA-CAAHR muscle quantified via bulk RNA sequencing analysis (n=3 male mice per group). Analysis in panels A-E were performed using two-way ANOVA with Sidak’s post hoc testing for multiple comparisons. Error bars represent the standard deviation.

Next, immunoblotting for electron transport system proteins was performed to assess if/how AAV-HSA-CAAHR treatment impacted mitochondrial content/density (**Figure 9D,** all blot images shown in **Supplemental Figure 9**). In this regard, no significant main effects for treatment were observed (**Figure 9E**). However, significant main effects of sex were observed for UQCRC2, MTCO1, and NDUFB8 demonstrating that female mice had increased mitochondrial protein abundance, a result that likely reflects compensatory mitochondrial biogenesis in response to poor OXPHOS function seen in Figure 9A-C. In some cases (ATP5A and UQCRC2), females treated with AAV-HSA-CAAHR had significant higher protein levels compared to AAV-HSA-GFP treated mice. Considering that OXPHOS function, determined using respirometric analyses of isolated mitochondria, was impaired in AAV-HSA-CAAHR treated mice, the increase protein abundance of the electron transport system proteins ATP5A and UQCRC2 may be indicative a compensatory response to the enzymatic impairments observed in these mitochondria. RNA sequencing performed on total RNA isolated from the gastrocnemius muscle of AAV-HSA-CAAHR treated mice also demonstrated altered mitochondrial gene expression when compared to AAV-HSA-GFP treated mice (**Figure 9F**). For example, AAV-HSA-CAAHR increased mRNA expression levels for several genes involved in lipid metabolism (*Cpt2, Cpt1a, Acot1, Acsl3*), altered abundance of mRNA’s for genes related to pyruvate metabolism (*Pdk1, Pdk4, Pdp1*), the tricarboxylic acid cycle (*Cs, Idh2, Ogdh*), and lactate metabolism (*Ldhb*). Numerous OXPHOS related genes related to NADH dehydrogenase (*Ndufaf6, Ndufaf4, Ndufs4*) were downregulated, whereas several cytochrome c oxidase genes (*Cox17, Cox18, Cox6b1*) were upregulated. AAV-HSA-CAAHR treated muscles also displayed a significant induction of *Ucp3*, an uncoupler of the respiratory system; as well as *Polg*, an enzyme involved in mitochondrial DNA repair. In totality, these data establish a novel role of chronic AHR activation in modulating skeletal muscle mitochondrial energetics.

## DISCUSSION

In this study, we first identified the presence of significant AHR activation in the limb skeletal muscle of mice and human patients with CKD and PAD and confirmed that tryptophan-derived uremic metabolites activate the AHR pathway in mouse and human muscle cells *in vitro*. Skeletal-muscle specific ablation of the AHR in mice with CKD improved limb muscle perfusion recovery, increase arteriogenesis and paracrine vasculogenic signaling, attenuated muscle atrophy, and improved muscle contractile function and mitochondrial OXPHOS in an experimental model of PAD. Furthermore, viral-mediated skeletal muscle-specific expression of a constitutively active AHR in mice with normal kidney function exacerbated the ischemic myopathy evidenced by smaller muscle masses, reduced contractile function, histopathology, and lower mitochondrial OXPHOS function and respiratory capacity. Importantly, deletion of the AHR in skeletal muscle had no effect on muscle health or function in mice subjected to sham surgery or those without CKD where ligand abundance is low. Coupled with the results of a recent study ^21^, the findings herein suggest that future studies should explore the therapeutic potential of AHR inhibition as an adjuvant treatment option for PAD patients with CKD.

The pathogenesis of PAD unquestionably stems from the development of atherosclerosis in the peripheral vasculature resulting in compromised blood flow to the lower limbs. However, other factors beyond atherosclerosis have emerged as important contributors to limb symptoms and adverse limb events including microvascular disease ^61,62^, endothelial dysfunction ^63^, and skeletal muscle dysfunction ^64,65^. Notably, CKD is recognized to be a strong, independent risk factor for PAD ^66–68^ which has also been demonstrated to negatively impact both the vasculature ^69,70^ and skeletal muscle independent of PAD ^71^. The accumulation of several toxic, protein-bound, uremic metabolites including indoxyl sulfate, indole-3-acetic acid, L-kynurenine, and kynurenic acid (among others) have been directly linked to vascular and skeletal muscle pathophysiology in CKD ^23,24,32,70,72,73^. Concerning for PAD patients, hazard ratios for the incidence of adverse limb events (after adjustment for age, sex, and eGFR) demonstrated a strong link to the levels of these specific uremic toxins ^21^. Interestingly, each these tryptophan derived uremic toxins are established ligands of the AHR, suggesting that this ligand-activated cytosolic receptor could be a common link between uremic toxicity and the muscle and vascular pathology in CKD. Mechanistically, AHR activation has been shown to promote endothelial cell senescence ^74^, promote atherosclerosis, inflammation ^36^, and thrombosis ^73^, as well as impair angiogenesis ^21,34,48^. Importantly, AHR global knockout mice display improved limb perfusion recovery following treatment with the AHR ligand benzo[*a*]pyrene following hindlimb ischemia^48,75^. An intriguing observation from the current study is that deletion of the AHR in skeletal muscle cells of mice with CKD and PAD improved limb perfusion recovery within the gastrocnemius muscle (**Figure 2G**). Improved perfusion recovery corresponded with increased arteriole density in both male and female AHR^mKO^ mice with CKD (**Figure 2K**), increased pericyte abundance in male AHR^mKO^ mice, but normal capillary density following FAL. Gene expression analysis also revealed increase expression of *Nos2*, a well-established vasodilatory gene, in AHR^mKO^ mice with CKD. Considering that AHR deletion occurred only in myofibers, these findings suggest that AHR activation alters paracrine signaling events between the skeletal muscle and vasculature within the ischemic limb. To this end, interrogation of intercellular communication processes via snRNA sequencing demonstrated that AHR^mKO^ mice with CKD had greater angiogenic signaling (*Vegf, Notch, Fgf*) between myonuclei and endothelial cells (**Figure 6**). qPCR confirmed higher expression levels of several vasculogenic genes in both the ischemic and non-ischemic limbs of AHR^mKO^ mice with CKD. A functional deficit in angiogenic growth factor secretion was further confirmed in C2C12 myotubes, firmly establishing an important role of AHR activation in disrupting muscle-vascular communication. In contrast to CKD mice, AAV-HSA-CAAHR treated mice displayed relatively normal gastrocnemius muscle perfusion recovery (**Figure 7D**), as well as normal capillary and pericyte levels (**Figure 7E,F**). Contributing factors to this discrepancy likely stem from the heterogenous infection of AAV to the hindlimb muscles and the superficial penetration depth of laser Doppler which does not measure into the depths of the gastrocnemius muscle. Moreover, laser Doppler flowmetry was performed using a small probe rather than a scanning imager which would have sampled more tissue area. However, it is worth highlighting that angiogenic signaling pathways and arteriole density were improved in AHR^mKO^ with CKD and decreased in AAV-HSA-CAAHR treated mice. Nonetheless, of great concern for the PAD patient, the findings in the current study and others ^21^ indicate that CKD negatively impacts both skeletal muscle and vasculature (endothelial cells, pericytes, and smooth muscle arterioles) health, two critical components regulating walking performance and limb outcomes.

Skeletal muscle mitochondrial alterations have been documented in both PAD patients ^39,40, 50– 55,76,77^ and CKD patients ^13,78–81^, and thus could serve as a site for the coalescence of these conditions in patients suffering from both conditions. In the current study, genetic deletion of the AHR in skeletal muscle significantly improved mitochondrial OXPHOS in the ischemic muscle from CKD mice (**Figure 4**). However, this effect was primarily driven by the enhanced OXPHOS function in male mice rather than female mice, especially when mitochondria were fueled by carbohydrates but not fatty acid sources. Conversely, AAV-driven AHR activation impaired mitochondrial OXPHOS in the ischemic limb of mice from both sexes when mitochondria were energized by a mixture of carbohydrates and fatty acids (**Figure 9**), suggesting that AHR mediates mitochondrial metabolism in a substrate-specific manner. Previous studies have identified sex-dependent transcriptional changes in the liver following treatment with the AHR ligand, TCDD, including numerous transcripts involved in metabolism ^82^. While further work is needed to identify the specific mechanisms underlying AHR-dependent mitochondrial dysfunction in CKD/PAD, similar observations have been made in other cell/tissue types. For example, AHR activation has been suggested to decrease mitochondrial membrane potential, impair respiration, and/or increase ROS in spermatozoa ^83^, hepatocytes ^84,85^, pancreas ^86^, cardiomyocytes ^87^, and the brain ^88^. Similarly, treatment of cultured myotubes with tobacco smoke condensate (which contains numerous AHR ligands) or expression of a CAAHR was reported to impair mitochondrial respiratory capacity ^38^.

The AHR is most known for its role as a ligand activated transcription factor involved in xenobiotic toxicity. However, several reports have linked AHR activation to ischemic tissue damage. For example, cerebral ischemia (i.e., stroke) induces AHR activation via L-kynurenine and subsequent deletion of the AHR attenuated ischemic brain damage in mice ^89^. This finding has been replicated using an AHR inhibitor in rats subjected to cerebral ischemia/reperfusion ^90^. In cardiomyocytes, activation of the AHR by kynurenine has been shown to drive ROS production and promote apoptosis which could be prevented by AHR inhibition ^91^. This study also demonstrated higher levels of cardiomyocyte apoptosis and larger infarct size following myocardial infarction in mice treated with exogenous kynurenine ^91^.

Paradoxically, it is worth noting that kynurenic acid, a known AHR ligand, has also been reported to have cardioprotective effects in the setting of ischemia/reperfusion, although this cardioprotection does not appear to require the AHR ^92^. Similarly, AHR activation in T-cells has also been reported to exert cardioprotective effects in mice with myocardial infarction ^93^. In the current study, analysis of cell populations via snRNA sequencing five days following the induction of limb ischemia demonstrated that deletion of the AHR in skeletal muscle preserved mature myonuclei (Type IIb and IIx) (**Figure 5**). Given the acute phase of ischemic injury at the time of muscle collection, this finding may be consistent with the prevention of cell death pathways that has been reported in ischemic cardiomyocytes treated with AHR inhibitors ^91^.

An unexpected finding of this study was the deletion of the AHR in skeletal muscle improved muscle size, muscle strength, and mitochondrial function in male, but not female mice with CKD. However, it is interesting to note that female AHR^mKO^ mice with CKD displayed improved perfusion recovery compared to their AHR^fl/fl^ counterparts with CKD (Figure 2), suggesting that the pathological effects of AHR activation in muscle may be disentangled from the limb hemodynamics. This assertion is support by previous work from our group demonstrating that muscle-targeted therapies in mice subjected to FAL can also improve muscle contractile function without corresponding improvements in limb perfusion recovery or angiogenesis^94^, as well as vascular-centric therapies improving angiogenesis without enhancing muscle contractile function^46^. While the exact mechanisms underlying the sex differences are unknown, several previous studies have reported sex-differences in the AHR reactivity in non-muscle tissues. For example, the hepatic transcriptional response to treatment with TCDD was muted in female mice when compared to males ^82^. In rodents, CKD has been shown to alter AHR abundance in the kidney in a sex-dependent manner^95^. Additionally, the severity of gastric tumors induced by expression of a constitutively active AHR was worse in male mice compared to female mice ^60^. Intriguingly, the AHR/ARNT heterodimer has been reported to be physically associated with the estrogen receptor and may regulate its transcriptional activity ^96^, providing a potential explanation for the sex-dependent effects observed in this study. The AHR has also been implicated to promote proteasomal degradation of the estrogen receptor via the cullin 4B ubiquitin ligase ^97^. Additional mechanistic experimentation is required to fully establish the sex-dependent AHR mechanisms in the context of CKD-PAD pathobiology.

There are some limitations of the experimental approaches used in this study that are worthy of discussion. First, this study employed younger mice despite the well-known association between PAD prevalence and increasing age. However, it is important to note that mice were enrolled into studies between the ages of 4-6 months to ensure the skeletal muscle and vasculature were fully matured. Further to this, PAD/CLTI patients with CKD often suffer from numerous other co-morbid conditions such as hypertension, diabetes, and hyperlipidemia that were not present in the mice used in this study. These comorbidities layered on top of the genetic and environmental risk factors (i.e., smoking, poor diet, lack of physical activity) that are typically present are all contributing factors to patient outcomes. Additionally, PAD/CLTI is a progressive disease where atherosclerosis develops slowly over time and is hastened by the presence of CKD. In contrast, a rapid and severe decrease in limb blood was induced by FAL in the experiments herein. As such, the conditions of chronic ischemia are not well modeled by the FAL surgery employed in this study. Nevertheless, a benefit of the FAL surgery stems from the ability to use clear anatomical landmarks for surgical ligatures which results in a consistent post-operative outcome that facilitates testing therapeutic/biological interventions, such as AHR deletion. Finally, it was not possible to perform temporal analyses within this study design which could have allowed exploration of the dynamic changes in arteriogenesis and angiogenesis in the ischemic hindlimb and the impact AHR activation had on these important processes.

## CONCLUSIONS

These data demonstrate that chronic *Ahr* activation, such as that in chronic kidney disease, is an important regulator of murine ischemic myopathy. *Ahr* activation, which has been previously observed in blood from PAD patients with CKD and linked to adverse limb events ^21^, was also observed in skeletal muscle specimens from PAD patients with CKD. Ablation of the AHR in skeletal muscle of mice with CKD significantly improved limb perfusion recovery and preserved paracrine vasculogenic signaling, improved muscle mass and contractile function, as well as enhanced mitochondrial energetics in an experimental model of PAD. Overall, these findings are an important step in elucidating the complex pathobiology of PAD in the context of renal insufficiency and provide additional support for testing of interventions that diminish *Ahr* signaling in these conditions.

## ABBREVIATIONS

AAV: adeno-associated virus
AHR: aryl hydrocarbon receptor
AHR^mKO^: muscle-specific AHR knockout mouse
CAAHR: constitutively active AHR
CKD: chronic kidney disease
FAL: femoral artery ligation
GFP: green fluorescent protein
GFR: glomerular filtration rate
MuSC: muscle stem cell
OXPHOS: oxidative phosphorylation
PAD: peripheral arterial disease
ROS: reactive oxygen species

